# Deep learning predicts tissue outcomes in retinal organoids

**DOI:** 10.1101/2025.02.19.639061

**Authors:** Cassian Afting, Norin Bhatti, Christina Schlagheck, Encarnación Sánchez Salvador, Laura Herrera-Astorga, Rashi Agarwal, Risa Suzuki, Nicolaj Hackert, Hanns-Martin Lorenz, Lucie Zilova, Joachim Wittbrodt, Tarik Exner

## Abstract

Retinal organoids have become important models for studying development and disease, yet stochastic heterogeneity in the formation of cell types, tissues, and phenotypes remains a major challenge. This limits our ability to precisely experimentally address the early developmental trajectories towards these outcomes. Here, we utilize deep learning to predict the differentiation path and resulting tissues in retinal organoids well before they become visually discernible. Our approach effectively bypasses the challenge of organoid-related heterogeneity in tissue formation. For this, we acquired a high-resolution time-lapse imaging dataset comprising about 1,000 organoids and over 100,000 images enabling precise temporal tracking of organoid development. By combining expert annotations with advanced image analysis of organoid morphology, we characterized the heterogeneity of the retinal pigmented epithelium (RPE) and lens tissues, as well as global organoid morphologies over time. Using this training set, our deep learning approach accurately predicts the emergence and size of RPE and lens tissue formation on an organoid-by-organoid basis at early developmental stages, refining our understanding of when early lineage decisions are made. This approach advances knowledge of tissue and phenotype decision-making in organoid development and can inform the design of similar predictive platforms for other organoid systems, paving the way for more standardized and reproducible organoid research. Finally, it provides a direct focus on early developmental time points for in-depth molecular analyses, alleviated from confounding effects of heterogeneity.

## Introduction

Retinal organoids (RO), derived from embryonic and (induced-) pluripotent stem cells of human (1), mice (2) and fish (3), have become vital models for understanding the retina in development and disease. Yet, notable challenges remain and one key challenge, that is found in RO as well as in organoid systems generally, is their intra-and inter-organoid heterogeneity, which is introduced from the very onset of their assembly as well as throughout their life cycle. This heterogeneity is believed to be caused by, among others, the stochastic nature of differentiation processes and self-organization (4, 5). Organoids subjected to identical conditions within and across experiments often exhibit markedly different outcomes, including starkly contrasting and even opposing phenotypes (6, 7). RO, specifically, may vary in tissue outcomes such as the emergence of the functionally highly relevant retinal pigmented epithelium (RPE), their cell type diversity as well as more general phenotypic outcomes like symmetry, size and shape (8).

Some degree of variability and heterogeneity in organoids might be desirable, since it reflects the complexity of biological systems (9). However, unintended heterogeneity essentially prohibits unconfounded studies of early development and related phenotypes in organoids. Under those conditions it is impossible to predict whether a particular phenotype will emerge, and destructive analysis would preclude subsequent verification.

There is considerable interest in the (retinal) organoid community to conduct molecular analyses at single-cell resolution on organoids of different developmental stages to unveil their specific cellular composition and characteristics of developmental trajectories (10, 11). These approaches have even been combined in a multimodal manner with immunofluorescence imaging, profiling RO heterogeneity spatio-temporally within and across samples (12). *A priori* knowledge about whether a specific phenotype will emerge in any specific organoid within an experiment would consequently enable these molecular analyses to be purged from the confounding factor of heterogeneity.

Further, it is crucial to determine at which specific time windows a trajectory towards a specific phenotypic outcome is set during organoid development. Identifying these fate-defining windows results in a better understanding of the system and allows targeted efforts in improving culture conditions. Importantly, it will guide researchers to identify promising temporally matching targets specifically triggering the developmental routes for their tissue outcomes of interest (TOI). Ultimately, this would facilitate more consistency in studying these phenotypic outcomes, thereby enhancing the utility of organoid systems as organ and tissue models in development and disease.

Recent advances in deep learning (DL) technology allowed successful application of DL models to accurately segment organoids from images and analyze their structural features (13–16). Further, one study effectively trained a neural network to predict mRNA expression levels in relation to tissue differentiation in kidney organoids (17), while another one successfully predicted the spatial distribution of marker genes within retinal organoids (18), demonstrating the potential of DL to bridge imaging and molecular data. A convolutional neural network (CNN) was successfully employed to recognize the presence of retinal precursor structures in early retinal mouse organoids to predict later retinal identity, further underscoring the utility of these techniques in developmental biology (19). When it comes to supervised DL in organoid research, the acquisition of annotated high-quality datasets spanning up to hundreds of thousands of images has been recognized as one of the major limitations in this area (20). In mouse and human RO, it typically takes weeks and months, respectively, until first signs of differentiated retinal cell types can be detected (2, 10). This creates a substantial time investment when used for proof-of-concept studies, where experimental turnovers are typically high and hard to anticipate. Therefore fast-developing, non-mammalian, yet vertebrate organoid systems, like those derived from fish, that recapitulate all key developmental aspects of mammalian organ development are particularly suited for these types of studies (3, 21).

Overall, despite these advances in DL powered organoid analysis and the promises DL holds when it comes to recognition of image patterns beyond human capabilities (20), there remains a notable gap in the application of DL to predict whether and to what extent tissues and phenotypes will develop within organoids.

In this proof-of-concept study, we demonstrate how DL can predict tissue outcomes in medaka fish RO well before these outcomes visibly emerge. To achieve this, we acquired a time-lapse bright-field image data set spanning 988 organoids and 117,249 images. Using expert annotation and advanced image analysis of organoid morphology, we characterize the inter-and intra-RO heterogeneity for two TOI, namely RPE and the lens, as well as for the organoids’ global morphology over time. Finally, we utilize DL to break through this heterogeneity and accurately predict both the emergence as well as the sizes of RPE and lens tissue formation at very early developmental time points on an organoid-by-organoid basis. This significantly refines our understanding of the timeline when these tissue outcomes are determined during RO development. Our approach shows great potential for advancing our understanding of tissue and phenotype decision-making in organoid development across types and species. Moreover, it enables access to early developmental time points for in-depth molecular analyses, free from the confounding effects of tissue outcome and phenotype specific inter-organoid heterogeneity.

## Results

### Retinal organoids display inter-and intra-experimental heterogeneity with specific tissue outcomes

In order to show that deep learning (DL) can predict tissue outcomes in retinal organoids (RO) well before they emerge, we first generated a comprehensive dataset that would enable us to monitor tissue development at high temporal resolution. For this, single RO derived from embryonic pluripotent cells of *Oryzias latipes* were seeded in 96-well plates and high throughput time-lapse imaged every 30 minutes for a total duration of 72 h using an automated widefield microscopy platform (**Figure 1A** and **Methods**). This way, we were able to assemble a dataset totaling 988 organoids with 117,249 images. As specific tissue outcomes of interest (TOI), we chose retinal pigmented epithelium (RPE) and the formation of lenses. Both tissues are highly physiologically and developmentally relevant ((8, 22), Zilova, L. and Wittbrodt, J., unpublished) in the organoid context as well as easily observable in bright-field imaging. While RPE formation was triggered on-demand by the addition of the recombinant WNT surrogate-Fc Fusion Protein (Wnt-surrogate) (21) and therefore served as an example for an induced tissue formation, lenses were found to develop independently of Wnt-surrogate supplementation to the media (**Supplementary Figures S1A**, **S1B**). Thus, they served as an example of a spontaneously emerging TOI in the organoids.

**Figure 1:**
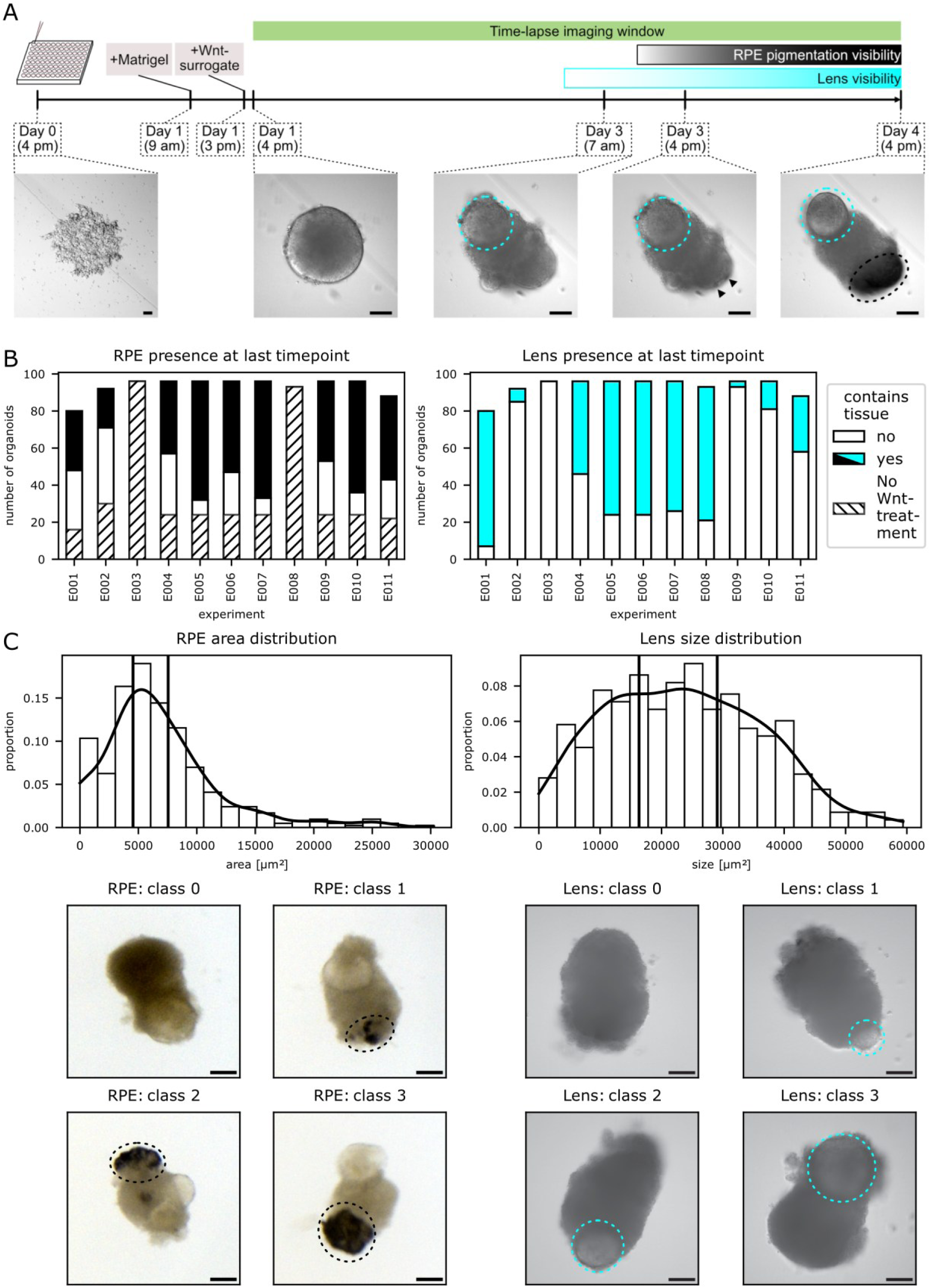
Tissue-specific heterogeneity of retinal organoids across development. **A** Experimental overview. Retinal organoids derived from *Oryzias latipes* embryonic pluripotent cells were treated with Wnt-surrogate 23 h after seeding and subsequently imaged by time-lapse widefield microscopy for 72 h every 30 min. The development of two tissue outcomes of interest (TOI) were followed - the lens (cyan circle) and the retinal pigmented epithelium (RPE; marked by arrowheads and a black circle) - which begin to visibly emerge at 59 h and 68 h of organoid development, respectively. Scale bars: 100 µm **B** Quantification of the development of the TOI. Organoids were classified for the presence or absence of RPE (left graph) or the formation of lenses (right graph) at 96 h of organoid development. Although culture conditions were kept uniform across experiments, there was high heterogeneity regarding the number of organoids developing the TOI. In the left panel (RPE emergence), the proportion of organoids without Wnt-surrogate treatment is indicated by diagonal line pattern for clarity. Lens emergence was found to be independent of Wnt-surrogate treatment and treatment conditions are thus not indicated. **C** Quantification of RPE and lens areas. Histogram of RPE (top left) and lens (top right) areas per organoid across the dataset. Four classes of TOI sizes were defined based on the quantile distributions (vertical lines; see also Methods), with organoids without development of the TOI being classified as class 0 and thus not included in the histogram. Class 1, 2 and 3 were assigned for areas below the 33rd, 66th and 100th percentile, respectively. Example images are given for the respective TOIs (RPE: stereomicroscopy, Lens: time-lapse widefield microscopy). Scale bars: 100 µm.

To delineate and cross-validate the visibility of the respective tissues over time, we first assembled an expert panel of seven researchers with expertise in retinal organoid biology. The expert panel annotated a randomly selected, balanced subset of 4000 images, equally covering organoids with both the presence and absence of RPE or lenses (see **Supplementary Table S1** and **Methods**). While the emergence of RPE became apparent at approximately 44 h into the imaging window (68 h of organoid development; **Figure 1A**, **Supplementary Figure S1C**), lenses were confidently detectable approximately 35 h after the onset of image acquisition (59 h of organoid development; **Figure 1A**, **Supplementary Figure S1D**).

Next, we set a ground truth for the presence or absence of RPE and lenses, respectively. For this, all organoids from the whole dataset were classified independently by two experts from the expert panel at the last time point imaged (96 h of organoid development). The presence of RPE was determined using stereomicroscopy due to its increased sensitivity for RPE detection compared to the time-lapse imaging data, which was confirmed by the expert panel (**Figure 1A, Supplementary Figure S1C** and **Methods**). Lens formation was evaluated for each organoid from a z-stack of the time-lapse imaging data (**compare Figure 1A**).

We found a significant inter-and intra-experimental heterogeneity regarding tissue emergence of RPE and lenses in the RO, despite having taken extensive care to maintain constant and highly reproducible culture conditions. Even the on-demand induction of RPE by Wnt-surrogate did not ascertain the emergence of RPE in every given organoid (**Figure 1B, Supplementary Figure S1A, S1B** and **Supplementary Table S2**). We quantified the RPE and lens areas in the stereomicroscopy and time-lapse images, respectively, as an approximation for the RPE amount and lens size that developed in each organoid. Based on the distribution of the quantified areas, we then grouped each organoid into one of 4 classes (**Figure 1C**). Class 0 denoted the absence of tissue emergence in a given organoid, while class 1, 2 and 3 were assigned for areas below the 33rd, 66th and 100th percentile, respectively. As it was the case for RPE and lens emergence, RPE amount and lens sizes showed a similarly large heterogeneity within and across experiments (**Supplementary Figures S1E, S1F**). Moreover, increasing concentrations of Wnt-surrogate (1, 2 and 4 nM) applied did neither reliably ascertain the emergence nor increasing amounts of RPE to develop (**Supplementary Figure S1A** and **Supplementary Table S2**).

These results showed that the tissue formation and sizes of both RPE and lenses in each organoid are heterogeneous and could not be ascertained, even when induced in a controlled fashion via an external stimulus.

### Development of a large-scale time-lapse image analysis platform

For the purpose of analyzing the time-lapse images on a large scale, we next developed a python-based analysis pipeline, starting with a DL guided image segmentation. The segmented images were processed through our analysis platform, enabling us to quantify shape descriptors and image moments, among others, over time (total number of parameters: 165). This generated a descriptive, tabular dataset containing morphological characteristics of the organoids, termed morphometrics (**Figure 2A**). Distance measurements in conjunction with dimensionality reduction (**Figure 2B**, **Supplementary Figure S2A**) revealed that organoids were more similar to each other at early time points, but progressively diverged as development advanced. This suggested dynamic interindividual changes in their morphological characteristics over time (**Figure 2C**), which are in accordance with the literature reporting inter-and intra-RO heterogeneity on a transcriptional level over time (12). Although global morphological changes might partially be induced by the addition of Wnt-surrogate to the culture media, as has been previously reported in organoid systems for other Wnt agonists (23), the same trends were found in RO without addition of Wnt-surrogate (**Supplementary Figure S2B**). When we examined the morphological changes per RO over time, we could observe a considerably higher morphological change-rate in the second half of the imaging window compared to the first half, suggesting a higher organoid plasticity in later developmental stages (**Figure 2D**). This again was found in RO without addition of Wnt-surrogate as well (**Supplementary Figure S2C**). Therefore, heterogeneity was not only found in a tissue-specific manner but was also reflected globally in the temporal dynamic of the RO morphology.

**Figure 2:**
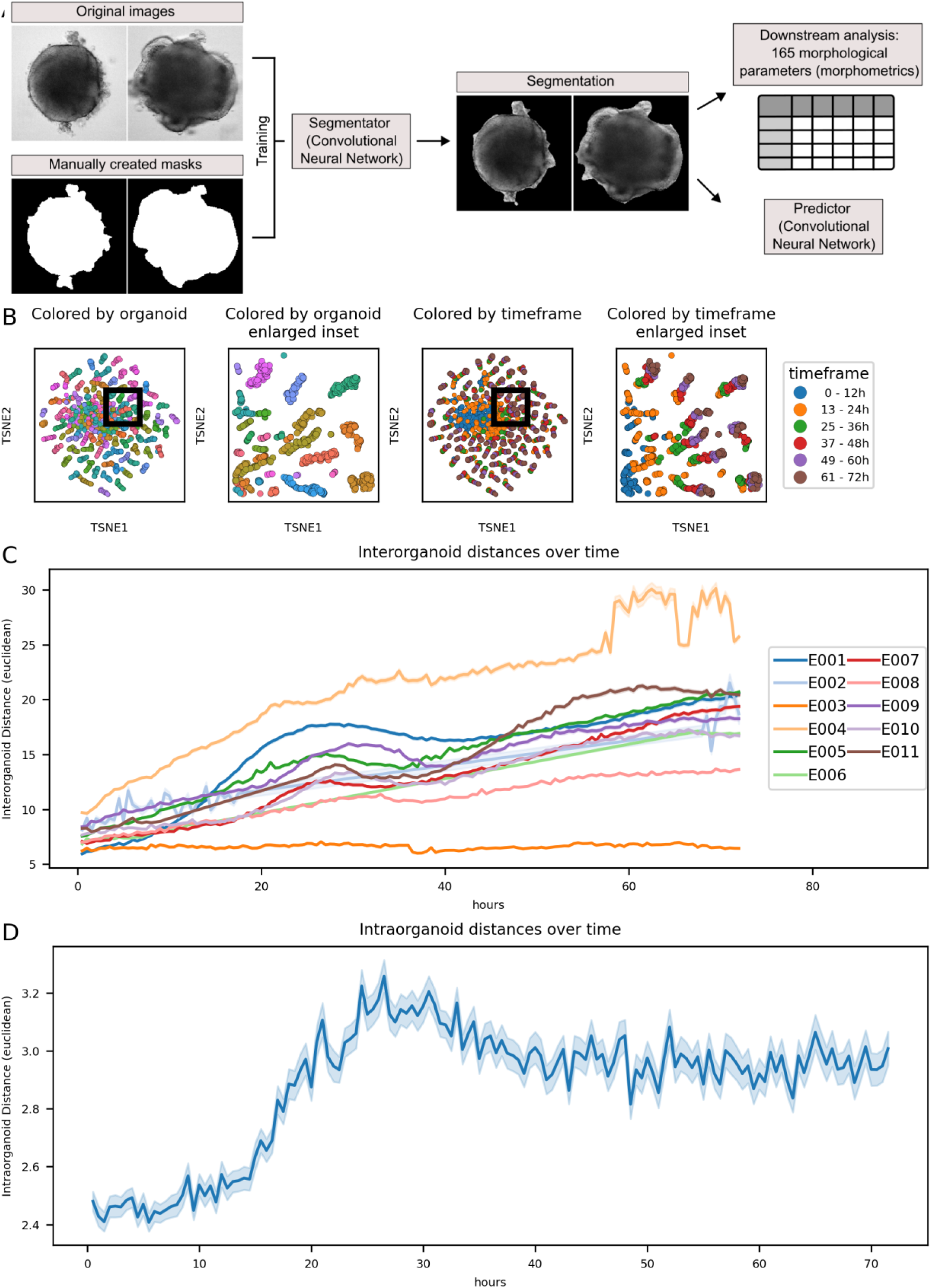
Global morphological heterogeneity of retinal organoids across development. **A** Image segmentation and analysis pipeline. A convolutional neural network (CNN) with the DeepLabV3 architecture was trained on 841 images and their manually annotated masks (left). The CNN was subsequently used to segment all images of the dataset that were finally subjected to a pipeline extracting a total of 165 morphological parameters (morphometrics, right), including, among others, shape descriptors and image moments. **B** Intra-experimental global morphological organoid heterogeneity. Time-series images from one representative experiment were analyzed using the image analysis pipeline and subjected to t-SNE dimensionality reduction. Data points were colored by individual organoids (left graph) and time frames of organoid development within the imaging window (right graph). While organoids clustered closely at earlier time points (up to 24 h), they strongly diverged at later time points, suggesting increasing inter-individual changes of their morphological characteristics over time. **C** Global morphological inter-organoid heterogeneity across experiments and time. Euclidean distances were calculated based on morphometrics data for each organoid and plotted as the mean pairwise distance between all organoids for each time point. Notably, inter-organoid distance increased over time in an experiment-specific manner, indicating an increasing morphological divergence of the individual organoids over time. **D** Intra-organoid morphological changes over time. Euclidean distances were calculated based on the morphometrics data for each organoid between time point *n* and time point *n+1*. The resulting metric reflects the amount of morphological changes during a time span of 30 min. While the relative changes are comparatively small at the beginning, the increase over time is suggesting more drastic morphological changes at later time points, consistent with the findings described in **B** and **C**.

### Deep learning predicts RPE and lens emergence well before visibility

Following the acquisition, annotation and analysis of our time-lapse RO imaging dataset, we next focused on the image classification task aimed at predicting the emergence of RPE and lenses.

As a reference for our prediction results, the expert panel was asked to *predict* the emergence of the tissues as well as their amount at the last time point of imaging using the same data subset as described above (**Figure 3A**, **Supplementary Table S1**). Importantly, to accurately reflect all the information they would have had during normal cell culture routines, the expert panel was given additional information about the age of each organoid in the images. Expectedly, the prediction accuracy of humans was low in the beginning and increased steadily towards later time points, consistent with the increasing visible emergence of RPE and lenses in the organoids (**Figure 3B, 3C**, compare **Supplementary Figure S1C, S1D**). Notably, the F1-metric plateaued at 0.7-0.8 for the expert evaluation at later time points, which is in line with our findings regarding the visibility of the respective tissues (compare **Supplementary Figure S1C, S1D**). Therefore, we concluded that an accurate *prediction* of tissue emergence of the TOI was infeasible for humans.

**Figure 3:**
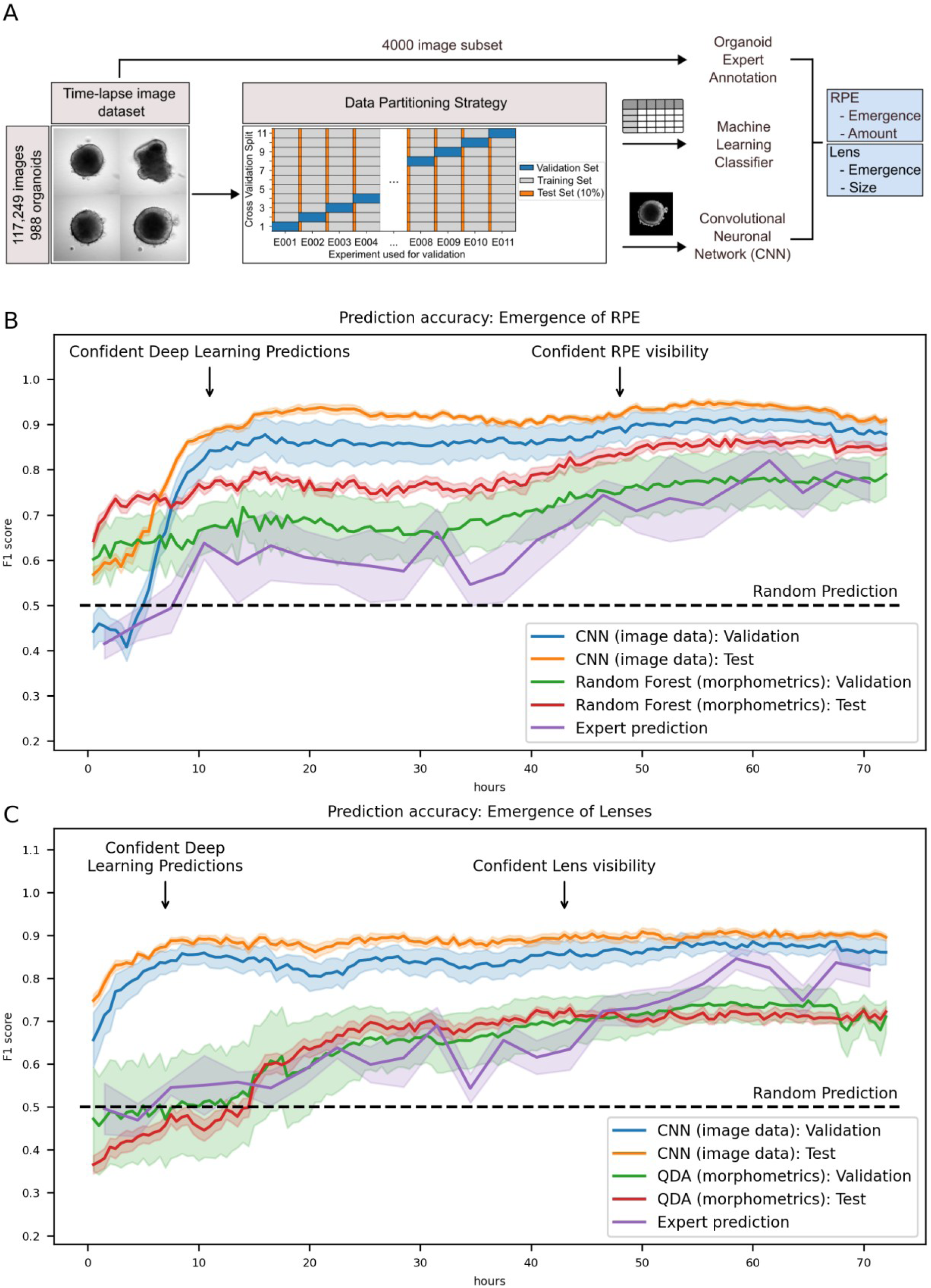
Deep-learning aided prediction of RPE and lens tissue emergence in retinal organoids. **A** Schematic representation of the data partitioning and machine learning model training strategy. Timelapse images were split into a training dataset and two distinct non-training datasets (data partitioning strategy): The test set was assembled from 10% of individual organoids that were imaged during the same experiments as the training set but were not trained on. The validation set was derived from organoid images acquired during a completely independent experiment that was not trained on. This strategy was repeated in a cross validation split 11-fold. The images were either analyzed via the image analysis pipeline or subjected directly after segmentation to a CNN ensemble. For the expert annotation, a randomly selected, balanced subset of 4000 images was designated as an evaluation set of which the experts chose *n* random images for classification and prediction (compare Supplementary Table S1 and Methods). **B/C** Prediction of RPE (B) and lens (C) emergence by deep learning. The indicated classifiers were trained on the segmented image data and morphometrics tabular data, respectively and evaluated on the test and validation set as described in A and Methods. While the classifiers trained on the tabular data could reproduce (C) and even slightly outperform (B) the human prediction accuracy, suggesting mostly tissue recognition, the CNN ensemble is able to predict RPE (B) and lens (C) emergence well before visibility at 11 h and 4.5 h, respectively (F1-score > 0.85). Lines represent the mean F1-score per time point, while shaded areas represent the standard error of the mean (SEM) over all test-experiments, validation-experiments and human evaluators, respectively. Horizontal dotted lines highlight the hypothetical accuracy of random predictions. QDA: Quadratic Discriminant Analysis

For machine-learning model training, we deliberately created two sets of test data that were not used for training. The first set, which we termed test set, was derived from 10% of organoids that were imaged within the same experiments that were used for training. The second set, termed validation set, contained organoids that were imaged in a completely independent experiment. Finally, we used a cross-validation strategy, where each of the acquired experiments was set as the validation experiment once and the training was performed on 90% of organoids of the remaining experiments (**Figure 3A**). By using this strategy, we were able to evaluate our model’s accuracy within as well as across the maximal breadth of inter-experimental variation in our data set.

In a first attempt for a machine-learning guided prediction of RPE and lens emergence, we aimed to predict the emergence of the respective tissues from tabular morphometrics data obtained from our image analysis pipeline. To facilitate this, we benchmarked several machine learning classifiers via conventional cross-validation in conjunction with hyperparameter tuning of selected classifiers to select the most suitable one for the image classification task (**Supplementary Figure S3A, S3B** and **Methods**). Finally, a Random Forest classifier and Quadratic Discriminant Analysis were chosen for the prediction of RPE and lens emergence, respectively.

RPE emergence was initially predicted at F1-score accuracies of 0.65-0.75 with this model. The accuracy slightly increased between 30-45 hours (42-63% of the imaging window) to a F1-score of 0.8, as time points approached the first visibility of the tissue. Notably, the accuracy was slightly outperforming the human prediction for all time points (**Figure 3B, Supplementary Figure S1C, S4A, S4B**). These results indicated that the classifier trained on tabular image analysis data reliably recognized tissue presence and was somewhat able to predict RPE emergence beforehand, yet only slightly outperforming human predictions. Although initially predicting lens emergence at random, we found a slight, but distinct increase in prediction accuracy to an F1-score of 0.6-0.7 with the model between 15-25 hours (21-35% of the imaging window). However, classification based on the morphometrics data was found to be inferior to the human prediction at later time points, suggesting that the morphometrics data did not adequately capture features associated with lens tissue emergence (**Figure 3C, Supplementary Figure S4C, S4D**).

In a second attempt, we trained an ensemble of convolutional neural networks (CNN) to classify images into two categories, predicting the final presence or absence of RPE and lenses at the last time point. The CNN demonstrated stable improvements during training as judged by the increase of the F1 metric (**Supplementary Figure S5**).

Strikingly, the DL model accurately predicted RPE and lens formation even at very early developmental stages (**Figure 3B, 3C, Supplementary Figure S6**). For the prediction of RPE, the network achieved its first substantial accuracy with a F1-metric above 0.85 at a median of 11 h after the onset of imaging, marking a critical early threshold for accurate RPE prediction (also compare **Supplementary Figure S6A, S6B**). The F1-metric was close to 0.9 for most of the remaining time points, providing far superior results compared to both the human prediction and the classification using tabular image analysis data (**Figure 3B**). These results indicated that the CNN was able to predict the emergence of RPE long before visibility of said tissue (compare **Supplementary Figure S1C**, **Figure 1A**) and even outperformed humans in organoids at time points when RPE was visible and detectable in stereomicroscopy but not detectable in the focus plane of the time-lapse images. For the prediction of lenses, the CNN ensemble was able to achieve its first confident prediction with an F1-metric above 0.85 at even earlier time points compared to the prediction of RPE (4.5 h). With a stable F1-score at that level over the remaining time points, the neural net ensemble clearly outperformed both the prediction capabilities of the experts and the tabular image analysis data-based model (**Figure 3C, Supplementary Figure S6C, S6D**). These results indicated that the prediction of lens emergence could be facilitated at very early developmental time points as well, even earlier than those of RPE using the same data set basis.

When we compared the performance on the test and validation sets as described above, we noticed that the models would generally perform more accurately on the test organoids, which were derived from the same experiments as the training set, compared to the validation organoids which were acquired in an independent experiment (**Figure 3B, 3C**, **Supplementary Figure S4, S6**). Additionally, we observed a substantial variability in the prediction accuracy in some validation experiments (**Supplementary Figure S4, S6**), underscoring the inter-experimental heterogeneity of organoid development and the necessity for our cross-validation strategy (**Figure 3A**).

### Tissue size prediction by deep learning

Next, we repeated our analyses attempting the prediction of the area of the RPE and lenses. As described above, we discretized the area obtained from stereomicroscopy and time-lapse image data into 4 classes based on the area distribution over all conducted experiments (**Figure 1C, Supplementary Table S2**).

We followed the same steps as above, including the human reference prediction, the classifier benchmark and hyperparameter tuning (**Supplementary Figure S7**) for the prediction from tabular image analysis data. The accuracy of the human prediction of tissue sizes increased steadily over the course of organoid development, consistent with findings obtained from the tissue emergence experiments. Human prediction reached F1-scores of 0.3-0.4 early on with a maximum F1-score of ∼0.53 for RPE areas (**Figure 4A**) and ∼0.65 for lens sizes (**Figure 4B**) towards the end of the imaging window after the onset of tissue visibility. This suggests that prediction of the amount of tissue is largely infeasible for humans.

**Figure 4:**
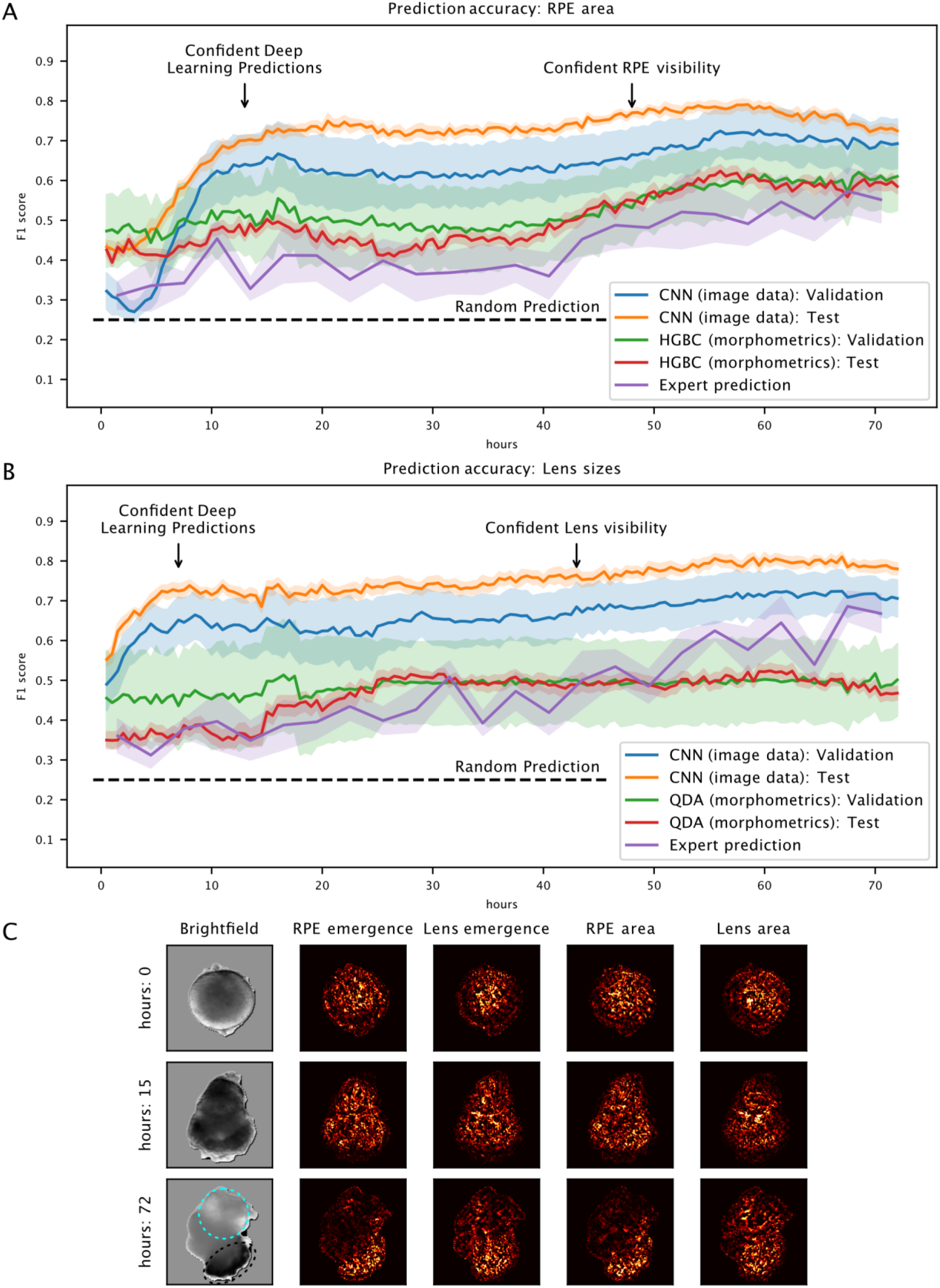
Deep-learning aided prediction of RPE and lens tissue sizes in retinal organoids. A/B. Prediction of RPE (A) and lens (B) tissue sizes by deep learning well before visibility. The indicated classifiers were trained on the segmented image data and morphometrics tabular data, respectively and evaluated on the test and validation set as described in Figure 3A and Methods. While the classifiers trained on the tabular data could reproduce or slightly outperform the human prediction accuracy, the CNN ensemble is able to predict RPE (B) and lens (C) tissue sizes at substantially higher accuracies well before visibility. Lines represent the mean F1-score per time point, while shaded areas represent the standard error of the mean (SEM) over all test-experiments, validation-experiments and human evaluators, respectively. Horizontal dotted lines highlight the hypothetical accuracy of random predictions. HGBC: HistGradientBoostingClassifier, QDA: Quadratic Discriminant Analysis **C** Relevance backpropagation. Pixels relevant to the CNN for the prediction of the indicated TOI were quantified using the GradientShap algorithm and plotted as absolute scaled values clipped to the 99.5th percentile. Dotted cyan line indicates the outline of the lens, dotted black line indicates RPE.

We next trained a HistGradientBoostingclassifier and a Quadratic Discriminant Analysis for the prediction of RPE and lenses, respectively, and observed a similar steady increase of prediction accuracy as judged by the F1-metric compared to the human prediction. The F1-score reached a plateau which was found to be slightly above the human performance for RPE at all time points and slightly inferior for lenses at later time points only (**Figure 4, Supplementary Figure S8**).

When we trained a CNN ensemble (**Supplementary Figure S9**), we observed a spike in prediction accuracy at very early developmental time points in line with our results obtained from the RPE emergence (compare **Figure 3B**) and the lens emergence (compare **Figure 3C**) with a final F1 metric of >0.7 for both tissues. Prediction accuracies were again stable after the initial spike, as it was the case for RPE and lens emergence predictions (**Figure 4 and Supplementary Figure S10**). Thus, for most RO the relative amount of RPE and the relative size of lenses could confidently be predicted around the same time points as their emergence, yet with lower overall accuracy. As it was the case for the RPE and lens emergence, we again noticed variance in the prediction accuracies between independent experiments (**Supplementary Figure S8, S10**).

Although both accurate emergence and size predictions of the TOI were acquired from training models with images from about 1,000 organoids, we suspect that as little as ∼500 organoids may be sufficient to reproduce the findings as reported here (**Supplementary Figure S11**).

Finally, we sought to find relevant structural information in the images that would guide the DL model’s decisions when predicting TOI at early time points before visibility. Using relevance backpropagation, we first found that pixels corresponding to visible RPE were highly relevant for the neural network to predict the presence and area of RPE (**Figure 4C**). Interestingly, the corresponding region of the organoid that would later contain RPE was not found to be highly relevant at earlier time points, indicating that relevant structural information is not confined to this region at all times and that the CNN does not identify the area where tissue will develop beforehand. For the prediction of lens emergence and its relative size, we obtained similar results but noted that the region that would ultimately contain the lens was neither highly relevant at earlier nor later time points (**Figure 4C**). Overall, we were not able to identify comprehensible organoid features with relevance backpropagation that would explain the decision making of the CNN and give further unequivocal insights into early indicators of organoid morphology for the development of the TOI.

## Discussion

### Model system heterogeneity as a challenge

In our study both non-spontaneous (RPE) and spontaneous (lens) tissue emergence, as well as their final tissue sizes, were found to vary between organoids and experiments undergoing the same differentiation protocol. Despite removing technical variability to the best of our ability, this type of heterogeneity seemed to be a rather inherent characteristic of our model system. Researchers are in general frequently confronted with considerable heterogeneity within their organoid model systems across and within experiments. These include variation in terms of cell type diversity and patterning, reaction to external stimuli and organoid morphology, among others (4, 24). The reasons for these heterogeneities are largely unknown, but are hypothesized to include, among others, deviations between differentiation protocols, batch-to-batch differences of cell culture material and cell sources (24–27). In line with this, we extend previous work (12) by demonstrating the *morphological* heterogeneity of RO within and across experiments over time using advanced image analysis tools. We comprehensively showed by distance analysis and dimensionality reduction that our model system exhibited intra-and inter-experimental variation of organoid morphology that consistently increased over time in an experiment-specific manner. While researchers in general undergo sincere efforts to minimize technical variation that would cause heterogeneity, there is yet no reproducible way to gain full control over all aspects of these complex model systems.

Due to the inherent intra-and inter-experimental heterogeneity of organoids, one can only be certain that a specific organoid possesses one TOI once it is properly and detectably established. Prior to this point, it remains uncertain whether a given organoid will develop the TOI. We refer to this period as the *Latent Determination Horizon* (**Figure 5**), representing the theoretical time window during which the decision toward developing the TOI is made in an organoid. Early detectable signs - such as specific transcriptional landscapes or single cells adopting the desired differentiation trajectory - may provide predictive hints but do not necessarily guarantee that the TOI will emerge. The stochastic nature of differentiation, along with unreliable spatio-temporal distribution of crucial cues (or the absence of such cues), may ultimately prevent the TOI from emerging. Moreover, some TOIs or desired phenotypes might not have any known early detectable markers.

**Figure 5:**
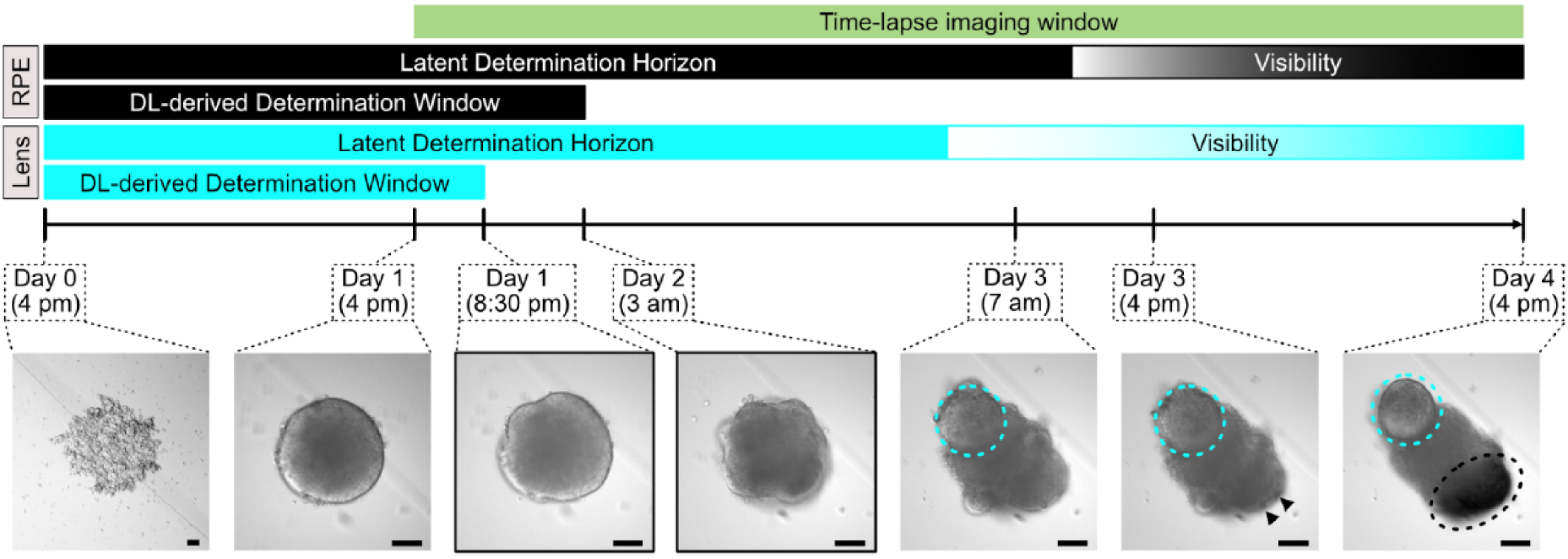
Deep learning mediated predictions of tissue outcomes narrow down phenotype determination windows in organoid development. Schematic representation of the retinal organoid developmental timeline showing the theoretical time windows (Latent Determination Horizon) during which the decisions towards RPE and lens tissue outcomes have to be made, and the narrowed-down time windows derived from deep learning (DL), where these decisions are actually being made. Lenses outlined in cyan and RPE indicated by arrowheads and a black circle. Scale bars: 100 µm.

### Comparison of machine learning approaches

To tackle the problem of organoid heterogeneity, we established a strategy that circumvents the challenge of heterogeneity within a larger group by predicting the outcome of a specific, singular organoid instead.

To accomplish that, we chose images as the fundamental, non-invasive data source, allowing studies of the respective organoids after analysis. This is in striking contrast to techniques like RNA sequencing that would require disintegration of the individual organoid and therefore prohibits studies at later time points of the same organoid. We built a high-temporal-resolution dataset of time-lapse images spanning about 1,000 organoids over the course of 72 h, which covered induction and development windows of RPE and lenses.

In order to predict tissue outcomes in a specific organoid, we chose two different approaches, on the one hand a “classical machine learning” (CML) approach based on tabular data obtained from our analysis pipeline and, on the other hand, a DL approach using the segmented images directly. When we compared the tissue outcome prediction accuracy of CML with that of the DL approach, DL consistently outperformed CML across all prediction tasks. Notably, CML did not perform significantly better than predictions made by human experts most of the time, indicating tissue recognition rather than prediction. This suggested that, for our prediction tasks, the relevant structural information within the images could not be effectively captured by existing bioimage analysis parameters or recognized by human experts through pattern recognition. In contrast, the predictive information within the time-lapse images is likely so complex and non-intuitive that only DL based on the segmented images was able to extract it successfully. Supporting this, our efforts to delineate the relevant structures for the DL classification at earlier time points were at best challenging, as we were not able to identify comprehensive organoid structures in the pixels the classifiers deemed relevant. This has been previously observed for relevance backpropagations in other contexts (28, 29). Taken together, we were able to reliably predict the emergence of RPE and lenses in individual organoids well before their actual visibility using DL.

### Deriving decision-making windows from prediction of tissue outcomes

By utilizing DL to predict tissue outcomes in organoids well before they visibly emerge, we could infer the time points in organoid development when the tissue outcome is already determined and identify those individual organoids that will adopt the TOI. This approach allowed us to narrow down the Latent Determination Horizons to *deep learning-derived determination windows* (**Figure 5**) - specific periods during which the decision toward the TOI in a given organoid is actually made. Consequently, efforts to decrease the variability of the TOI - such as modifying cell culture conditions - should be focused on these critical time frames of organoid development. However, it remains unclear whether the earliest time point at which DL can accurately predict the TOI corresponds to the actual biological decision-making moment or if it reflects a technical limitation of the model due to insufficient or inadequate training data. Therefore, the period of organoid development preceding this point should also be considered as temporal window during which decision-making occurs.

Using this methodology, we were able to substantially narrow down the decision-making window for both the emergence and ultimately the tissue sizes of RPE and lenses in RO. Interestingly, for both TOIs, these decisions coincide temporally to a large degree.

### Molecular analyses unconfounded by tissue-specific heterogeneity

One potential application of our analysis is the invasive study of tissue emergence in organoids. Consider a histological or transcriptomic study of organoids that will contain a specific tissue compared to organoids without it. As tissues have not yet emerged, grouping of the organoids can only be done at random for prospectively forming tissues. In conjunction with the invasive nature of the analysis, there is potentially no way of retrospectively identifying organoids, which would have developed the TOI and thus essentially prohibit such analyses. In line with our findings, even assigning the groups based on the on-demand induction of TOI by external stimuli would result in potentially highly inhomogeneous datasets with a strong confounding uncertainty of tissue outcome. Applying our model to predict TOI in organoids at time points where the models have shown confident prediction accuracies for the TOI will therefore significantly increase the homogeneity of the respective groups.

We deliberately selected an experimental setup that allows to make assumptions about the applicability of our models across experiments. This is especially important considering the large inter-experimental heterogeneities that were found in our dataset which we expect to extrapolate to other organoid systems. We observed a strong variability of model performance in independent experiments that would partially differ significantly from the accuracy observed from organoids that were imaged during the same experiments that were used for training (i.e. test set). Despite this variability, we are still confident that our technique can lead to a much higher purity of organoid groups compared to random group assignment.

### Applicability to other model systems and tissue outcomes

We strongly believe that our proof-of-concept study will pave the way for similar analyses across other model systems, tissue outcomes and more. In order to be applicable to other scenarios, the ground-truth annotation is of high importance. Even though the annotation of dark-pigment containing RPE and relatively large lenses might seem trivial, the presence of RPE and lenses were disagreed upon by the two independently annotating experts in a considerable fraction of organoids (3.9 % for RPE, 2.9% for lenses). This is an example for potential challenges that may ultimately require extensive work beforehand to ensure a highly accurate ground truth annotation. However, our study demonstrates that the effort is worth it, even if and in particular when the development of the organoids takes extended periods of time.

In this work, we used 1,000 organoids in total, to achieve the reported prediction accuracies. Yet, we suspect that as little as ∼500 organoids are sufficient to reliably recapitulate our findings. Therefore, our approach is readily applicable to any organoid model systems of choice given the reported success rate with a limited dataset. However, even though the indicated number of individual organoids may seem small in comparison to conventional datasets in DL, we note that the required number of unique organoids may exceed feasibility for some extreme models.

Beyond our current DL implementation, potential enhancements of our strategy could include the combination of tabular and image data, creating a broader dataset. Furthermore, a beneficial expansion of a time-lapse brightfield dataset might be the additional acquisition of epifluorescence images for every organoid. Apart from the additional information that transgenic reporter lines might provide, organoid autofluorescence and its spatio-temporal distribution might unlock image information beyond those obtained from bright-field imaging.

In summary, we have demonstrated that tissue trajectories in retinal organoids are reliably predicted well before the tissues visibly emerge. We achieve this by applying a deep learning approach to time-lapse bright-field image datasets, thus delivering prediction results instantaneously. This predictive capability provides vital insights into the timelines of decision-making during organoid development. Our deep learning and data curation framework can inform the design of similar frameworks for organoids across different types and species. This approach not only unlocks these developmental insights but also crucially grants access to early developmental time points for complementary in-depth molecular analyses, which were so far confounded by heterogeneity related to the tissue outcomes of interest.

## Methods

### Fish husbandry and maintenance

Medaka fish (*Oryzias latipes*) stocks were maintained according to the local animal welfare standards (Tierschutzgesetz §11, Abs. 1, Nr. 1, husbandry permit AZ35-9185.64/BH, line generation permit number 35–9185.81/G-145/15 Wittbrodt). The fish are kept as closed stocks in constantly recirculating systems at 28°C with a 14 h light/10 h dark cycle. For this study the following medaka strain was used: Cab strain (30).

### Generation of retinal organoids from medaka embryonic pluripotent cells

Retinal organoids were generated from medaka embryonic pluripotent cells as previously described (3) with some changes to the differentiation protocol. Briefly, blastula-stage (6 hours post fertilization; (31)) medaka embryos were taken as a source for primary embryonic pluripotent cells, which were resuspended in modified differentiation media (DMEM/F12 ((Dulbecco’s Modified Eagle Medium/Nutrient Mixture F-12), Gibco^TM^ Cat#: 21041025), 5% KSR (Gibco^TM^ Cat#: 10828028), 0.1 mM non-essential amino acids, 0.1 mM sodium pyruvate, 0.1 mM β-mercaptoethanol, 20 mM HEPES pH=7.4, 100 U/ml penicillin-streptomycin) following isolation. Cells were seeded in densities of approximately 1500 cells per organoid (approx. 15 cells/µl, 100 µl total) per well in low-binding, U-bottom shaped 96-well plates (Nunclon Sphera U-Shaped Bottom Microplate, Thermo Fisher Scientific Cat#: 174925) and incubated at 26°C without CO_2_ control. On day 1 at 9 am, organoids were washed with differentiation media, transferred into new low-binding, U-bottom shaped 96-well plates and Matrigel® (Corning, Cat#: 356230) was added to the media to a final concentration of 2% and a total media volume of 120 µl. Organoids were incubated for 6 h at 26°C under CO_2_ control. At 3 pm (day 1) WNT surrogate-Fc Fusion Protein (Wnt-surrogate; ImmunoPrecise Antibodies Ltd.; Cat#: N001, Lot: 7568) was added directly into the wells to final concentrations of 1 nM, 2 nM and 4 nM. Wnt-surrogate was stored in 250 nM aliquots in Wnt-surrogate dilution buffer at-20°C and pre-diluted to 100 nM in differentiation media prior to addition to the wells. Control organoids were left untreated. After Wnt-surrogate addition, organoids were subjected to time-lapse imaging. All experiments were performed by the same experimenter and times of experimental interventions were standardized to reduce technical variability in organoid generation.

### Organoid time-lapse imaging

Automated widefield time-lapse imaging of organoids was performed using the ACQUIFER Imaging Machine (ACQUIFER Imaging GmbH, Heidelberg, Germany) (32). Organoids were placed in 96-well plates and incubated at 26°C in the machine’s plate holder, without CO_2_ control. To prevent evaporation while enabling gas exchange, the plates were sealed with a Moisture Barrier Seal 96 (Azenta US, Inc.; Cat#: 4ti-0516/96). Brightfield images of each organoid were captured every 30 minutes over a 72-hour period (144 time points) using 70% illumination, a 20 ms exposure time, and a 10x objective (NA 0.3, CFI Plan Fluor10X). Images were acquired as z-stacks with five slices, spaced 50 µm apart, around the automatically determined focal plane, resulting in a total of 69,120 brightfield images per experimental replicate for 96 organoids. Z-stacks were used for expert annotation of tissue outcomes in the organoids, yet only the middle slice (slice 3/5) was used for image analysis and training. Following time-lapse imaging, organoids were imaged via stereomicroscopy (Nikon SMZ18 with a Digital Sight DS-Ri1 camera) to confirm RPE development.

Time-lapse recordings underwent manual quality checks, and wells exhibiting visible contamination were removed from further analysis and machine learning model training. Additionally, any time points where organoids were either out of frame or out of focus, as well as subsequent time points for the same organoid, were also excluded.

### Image ground truth annotation for machine learning model training

For training the machine learning models, time-lapse images of organoids were initially annotated for the presence of specific TOI in the organoids (RPE and lens) by two experts in medaka retinal organoid biology, working independently. Brightfield images, adjusted for contrast and brightness from the final time point in the automated widefield microscopy (time point 144th loop, 72 hours), were used for annotation for lenses, while stereomicroscopic images served as the gold standard for RPE detection and quantification (in concordance with the automated widefield microscopy). Annotations with consensus were assigned a high confidence score, while those with disagreement received a low confidence score. The decision on which expert annotation to include in the machine learning model training in case of a low confidence score was determined by a coin flip.

### Tissue outcome of interest quantification and classification

To quantify RPE areas, retinal organoids were first segmented in stereomicroscopic images using Yen’s global thresholding (33). Regions of interest (ROIs) were defined based on the segmented organoid areas and applied to the 8-bit converted raw images. Minimum-Maximum re-scaling of all pixel values within the ROI was used to enhance the contrast between RPE and non-RPE pixel values. Re-scaled images were then subjected to Sauvola’s local thresholding method, with a radius of 15 pixels, a k value of 0.5, and the recommended r value of 128 (34). The total area of all foreground pixels in the resulting binary images was then measured. For lens area quantification, lens areas were manually measured from slice 1, 3 or 5 of the acquired z-stack, depending on the focus plane of the lenses in the respective organoids, of the time-lapse brightfield images at the last time point (144th loop; 72 hours) using the oval tool in ImageJ (35). All lens areas were measured as circles. Since lenses in medaka retinal organoids are already transparent by day 4, their areas were assumed to correspond to the central cross-section of the mostly spherical lenses. This allowed for reliable measurements when comparing between samples, even if the lenses were positioned differently within the organoids.

Quantitative image analysis was used to classify organoids into four groups for machine learning model training. Group 0 included organoids without development of the TOI, while groups 1, 2, and 3 represented low, medium, and high levels of tissue sizes, respectively. Cut-off values for these groups were determined by calculating the 33^rd^ and 66^th^ percentiles of phenotype measurements from the entire training dataset. For RPE amounts, group 1 (low) included organoids with RPE areas below 4,541.73 µm^2^, group 2 (medium) had RPE areas between 4,541.73 µm^2^ and 7,548.51 µm^2^, and group 3 (high) consisted of organoids with RPE areas above 7,548.51 µm^2^. A similar approach was applied to lens sizes, using cut-off values of 16,324.85 µm^2^ and 29,083.23 µm^2^. This classification method ensured that groups were objectively defined based on the actual distribution of quantitative tissue outcome data, resulting in equally sized groups for a balanced dataset for machine learning model training.

### Organoid segmentation for image analysis and machine learning model training

In order to train a convolutional neural network to segment organoids, 841 organoid images were masked manually using Fiji (35) (v. 2.14.0/1.54f). The images and corresponding masks were read and downsampled to 512×512 px utilizing the INTER_AREA method of the open-cv package (v. 4.10.0). The dataset was subsequently split into a training set (90%) and a test set (10%). A pre-trained version of DeepLabV3 with a ResNet101 backbone (36) was trained with an initial learning rate of 3×10^-4^ and a batch size of 16 images for 500 epochs using the BCEWithLogitsLoss in the pytorch (v. 2.3.1) implementation. The classifier state with the lowest loss value on the test set was saved and used for further segmentation tasks after manual confirmation of the accuracy on images not belonging to either the training or test set.

### Large-scale time-lapse Image analysis

To measure morphological property metrics (termed morphometrics) of the organoids, the original images were downsampled to 512×512 px using the INTER_AREA method of the open-cv package and the mask was created using the trained segmentation-model. The mask was subsequently upsampled using the INTER_LINEAR implementation of open-cv and converted to a binary image with a threshold of 0.5 of the maximum pixel value. The mask and original image were subjected to sklearns regionprops function with the addition of custom implementations for the quantification of blur and the ImageJ shape descriptors, among others. Images where more than one mask were predicted or images where the mask would exceed a total diameter of 360 px x 360 px were excluded. For full implementation details refer to the source code.

### Distance calculation and dimensionality reduction

Morphometrics were calculated as described above and Z-scaled. Pairwise euclidean distances between organoids were calculated using the pdist function of scipy (v. 1.14.0, (37)) for each experiment and imaging time point. To calculate the distances between time points, the euclidean distances were calculated using the cdist function of scipy and quantile capped to the 2.5th and 97.5th percentile for each time point before plotting. tSNE dimensionality reduction was calculated using the scikit-learn (v. 1.5.1) implementation with standard settings. For the calculation of distances involving only organoids without Wnt-surrogate treatment, the scaling was performed before data subsetting to preserve the numerical space.

### Organoid classification using tabular metrics

Morphometrics were calculated as described above, Z-scaled and subsequently scaled to range from 0 to 1 to avoid negative values. In a first benchmark, classifiers in their sklearn (v. 1.5.1) implementation with standard settings were trained using leave-one-out training by reserving the data of one experiment for validation. The validation data were scaled to the same range as the training data using the fitted scalers obtained from the training data. The classifiers were trained on 90% of organoids while 10% of organoids were reserved as an internal test set. This benchmark was performed for the classification of binary classification of RPE and lenses as well as for the quantification of the tissue sizes thereof (**Supplementary Figure S3 and S7**). Selected classifiers for each readout were subjected to hyperparameter tuning based on their performance in the initial benchmark and known susceptibility to hyperparameter tuning. Hyperparameter tuning was performed using a custom implementation of the RandomHalvingSearch of sklearn with a reduction of factor of 3, a minimum_resource parameter of 1000 and 5-fold cross validation on the full dataset as described above (**Supplementary Figure S3 and S7**). The best performing parameters were saved and were used for the final evaluation. For full implementation details refer to the source code.

### Organoid classification using convolutional neural networks

Images were segmented as described above. Subsequently, a bounding box of 360 x 360 px was cropped using the mask and the resulting image was downsampled to 224 x 224 px. The images were scaled between 0 and 1 and stored together with the accompanying annotations in a custom class that would allow easy data access, splitting and downsampling (for full implementation details refer to the source code). Training data were split in a ratio of 90:10, where the images of individual organoids served as groups, yielding a training-and test-set, respectively. The validation data derived from an independent experiment were read in separately. For the training of convolutional neural networks, the pytorch (v2.3.1) implementations were used. Data were augmented using the albumentations (v1.4.12) package including rotations, distortions, random crops, rescalings and custom intensity adjustment, among others. The final normalization was performed only on the organoid pixels using a custom algorithm and used the same normalization values that were used for the initial model pretraining. Three models (DenseNet121 (38) ResNet50 (39) and MobileNetV3 (40)) were used in a pretrained state. Learning rates were determined beforehand for each readout using the pytorch-lr-finder package (https://github.com/davidtvs/pytorch-lr-finder). As a loss function and optimizer, CrossEntropyLoss and Adam were used in their pytorch implementations, respectively. Class weights for the loss function were calculated based on the distribution of the labels in the training set. To account for overfit, a weight-decay of 1e-4 was used for non-batchnorm-layers for all models. Gradients were clipped to a maximum of 1.0. A learning rate scheduler was implemented that would decrease the learning rate by 50% with a patience of 7 epochs based on the F1-score on the test data. Models were trained for at least 30 epochs, and the best performing model, judged by the loss on the test set, was saved. For each epoch, the loss values on the respective data subsets as well as the F1-score were recorded for plotting. For evaluation, neural networks were instantiated and calibrated using temperature scaling (41), using a slightly modified implementation of one of the original authors (https://github.com/gpleiss/temperature_scaling). The models were used for evaluation and their predictions were weighted based on the best F1-score. The class with the highest output probability was then used as the predicted label and used for subsequent statistical analysis. For full implementation details refer to the source code.

### Relevance backpropagation

For relevance backpropagation, the GradientShap algorithm of the captum (v.0.7.0, (42)) was used using the segmented image and zero-pixels of the same shape as baselines, 0.001 stdevs and n_samples of 200. The trained DenseNet121 model for the respective readout was loaded and the attributions were calculated for each single image, of which the first layer was used for the final analysis. The absolute attribution values were normalized using captum and displayed in the figure. For full implementation details refer to the source code.

### Data visualization and statistical analyses

Plots were generated using the matplotlib (v. 3.9.1) and the seaborn (v. 0.13.2) libraries. For ANOVA calculation (**Supplementary Figure S1**), scipy was used. Sketches in Figure 1, Figure 2, Figure 3 and Figure 5 were drawn using Inkscape 1.2.2 and Affinity Designer 1.10.5. F1-scores were calculated using the sklearn implementation with weighted averaging.

## Data Availability Statement

The data that support the findings of this study will be available on HeiData, the Open Research Data institutional repository for Heidelberg University, upon final publication of the manuscript. Raw image data are available upon reasonable request.

## Code Availability Statement

The code to reproduce analyses and figures are publicly available at github (https://github.com/TarikExner/orgAInoid).

## Acknowledgements

C.A. and T.E. were partially funded by the Structured Doctoral programme (*Strukturiertes Doktorandenprogramm zum Erwerb des Dr. med. und Dr. rer. nat.*) of Heidelberg University. L.H.A. was partially funded by postgraduate grant programmes (Beca de Apoyo a docentes para estudios de Posgrados, nivel doctorado and Forschungsstipendien - Bi-national betreute Promotionen/Cotutelle, 2023/24) of the Comisión Académica de Posgrado, Universidad de la República, and the German Academic Exchange Service (57645446). This work was made possible through the funding from the Deutsche Forschungsgemeinschaft (DFG, German Research Foundation) through Excellence Cluster “3D Matter Made to Order” (EXC-2082/1-390761711), the Forschungsgruppe FOR2509, WI 1824/9-1 and the European Research Council Synergy Grant IndiGene (number 810172) to J.W.. The authors acknowledge support by the state of Baden-Württemberg through bwHPC and the German Research Foundation (DFG) through grant INST 35/1597-1 FUGG.

## Author Contributions Statement

Conceptualization: C.A., J.W., T.E.

Investigation: C.A., N.B., C.S., E.S.S., M.L.H., R.A., R.S., N.H., H.-M.L., L.Z., J.W., T.E.

Visualization: C.A., T.E. Funding acquisition: J.W., T.E.

Project administration: C.A., T.E. Supervision: J.W., T.E.

Writing - original draft: C.A., T.E.

