## Supporting Information for "Deep learning predicts tissue outcomes in retinal organoids"

#### Supplementary Table S1

**Supplementary Table S1:** Number of images annotated per expert from the 4000 image balanced subset for tissue visibility and prediction of outcomes.

| expert ID | n_images |
|---|---|
| Expert1 | 551 |
| Expert2 | 955 |
| Expert3 | 1501 |
| Expert4 | 513 |
| Expert5 | 1612 |
| Expert6 | 500 |
| Expert7 | 777 |

### Supplementary Table S2

**Supplementary Table S2:** Dataset metrics across experiments. sd: standard deviation.

| Experiment_ID | Number of images | Condition (nM) | Number of organoids at last time point | % RPE | % Lens | RPE area (mean +/- sd) | Lens area (mean +/- sd) | Number of organoids for RPE classes [0/1/2/3] | Number of organoids for lens classes [0/1/2/3] | Time points acquired |
|---|---|---|---|---|---|---|---|---|---|---|
| E001 | 11520 | 0/1/4 | 80 | 40.00 | 91.25 | 11448 +/- 4376 | 33231 +/- 13163 | 48/1/6/25 | 7/12/14/47 | 1-144 |
| E002 | 728 | 0/1/4 | 13 | 23.08 | 0.00 | 21947 +/- 7926 | N/A | 10/0/0/3 | 13/0/0/0 | 1-35, 136-144 |
| E003 | 13824 | 0 | 96 | 0.00 | 0.00 | N/A | N/A | 96/0/0/0 | 96/0/0/0 | 1-144 |
| E004 | 13248 | 0/1/2/4 | 92 | 41.30 | 53.26 | 5852 +/- 3275 | 16584 +/- 8653 | 54/13/17/8 | 43/25/20/4 | 1- 144 |
| E005 | 13561 | 0/1/2/4 | 94 | 67.02 | 75.53 | 7267 +/- 3389 | 30200 +/- 11497 | 31/12/21/30 | 23/11/21/39 | 1-144 |
| E006 | 3589 | 0/1/2/4 | 91 | 51.65 | 73.63 | 4993 +/- 3650 | 24590 +/- 10521 | 44/19/20/8 | 24/19/23/25 | 1-29, 135-144 |
| E007 | 13053 | 0/1/2/4 | 90 | 67.78 | 73.33 | 5032 +/- 2254 | 17360 +/- 7489 | 29/25/27/9 | 24/29/34/3 | 1-144 |
| E008 | 12744 | 0 | 88 | 0.00 | 76.14 | N/A | 27887 +/- 9166 | 88/0/0/0 | 21/7/30/30 | 1-144 |
| E009 | 13244 | 0/1/2/4 | 91 | 46.15 | 3.30 | 7793 +/- 4817 | 7739 +/- 5045 | 49/11/10/21 | 88/3/0/0 | 1-144 |
| E010 | 12960 | 0/1/2/4 | 90 | 63.33 | 16.67 | 4827 +/- 2667 | 5938 +/- 5070 | 33/25/23/9 | 75/14/1/0 | 1-144 |
| E011 | 8778 | 0/1/2/4 | 77 | 54.55 | 33.77 | 4984 +/- 3778 | 15738 +/- 11243 | 35/22/5/15 | 51/16/5/5 | 1-144 |

### Supplementary Figure S1

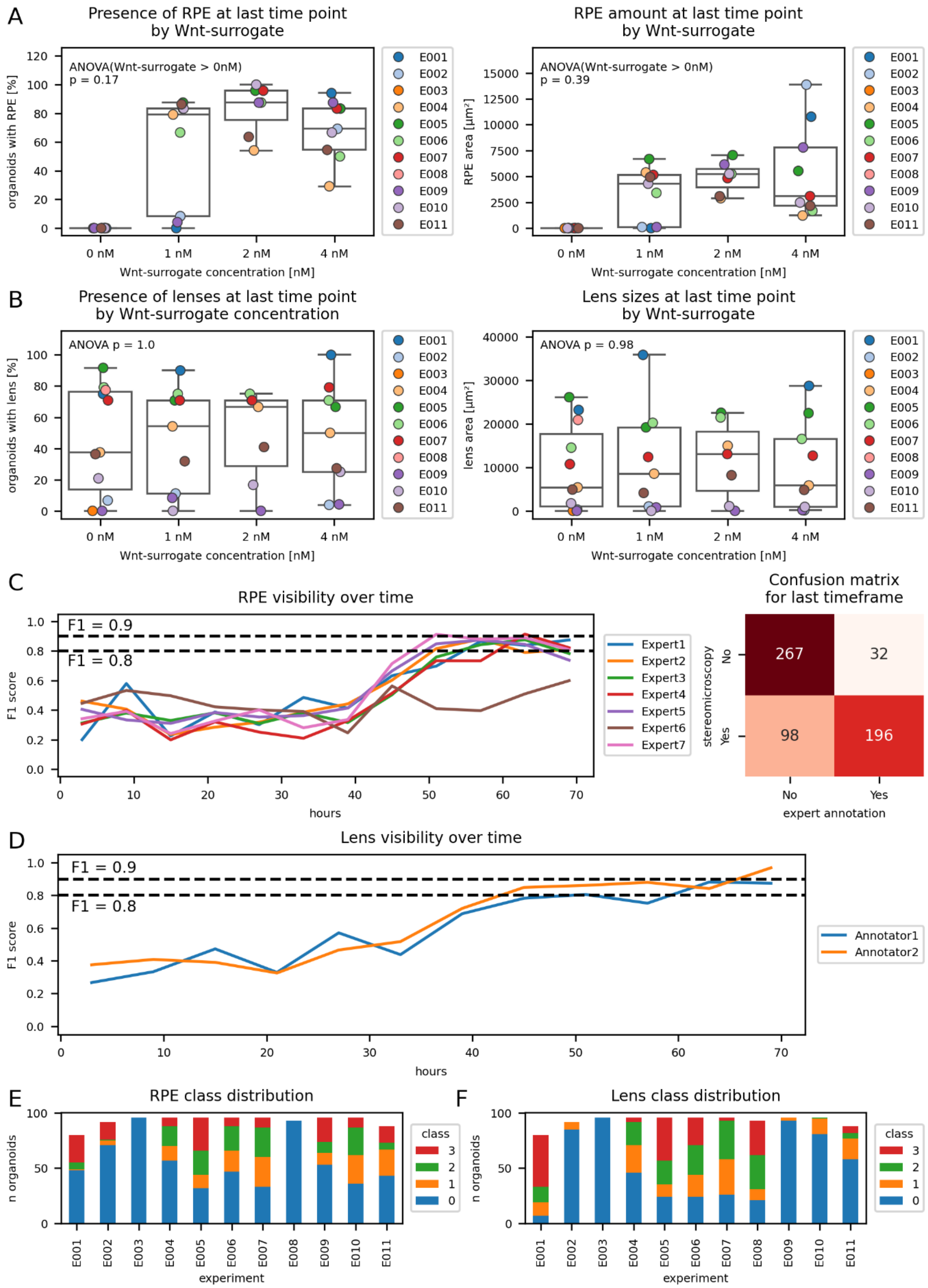

**Supplementary Figure S1: Tissue-specific heterogeneity and tissue visibility in retinal organoids across development with increasing Wnt-surrogate concentrations.**

**A** Development of retinal pigmented epithelium (RPE) and its amount by Wnt-surrogate concentration. Organoids were treated with the indicated concentrations of Wnt-surrogate and subjected to stereomicroscopy in order to detect the presence of RPE (left graph) and to measure its area (right graph). RPE development was observable after Wnt-surrogate treatment but absent in non-treated organoids over all conducted experiments. Notably, there was no consistent correlation of the concentration of the inducing agent and the rate (left graph) or amount (right graph) of RPE induction.

**B** Development of lenses and their sizes by Wnt-surrogate concentration. Organoids were treated with the indicated concentrations of Wnt-surrogate. Lenses were detected from the time-lapse widefield microscopy images (left graph) and the area was measured (right graph). Lens development and lens sizes were found to be independent of the Wnt-surrogate-treatment and -concentration. The indicated p-values were derived from a one-way ANOVA over all four groups. Data points are color coded for the respective experiments.

**C** RPE visibility over time and increased sensitivity of RPE detection by stereomicroscopy. Expert evaluation of the presence of RPE in each organoid over time (see Methods). F1-scores were calculated using the stereomicroscopy annotation as ground truth. RPE begins to visibly emerge around 68 h of organoid development. There is a notable difference between the stereomicroscopy derived ground truth and the human evaluation of the time-lapse images at later time points, suggesting an increased sensitivity of stereomicroscopy over the time-lapse images for the detection of visible RPE. The confusion matrix (right panel), summed over the last 6 hours of the imaging window, confirms the inferior sensitivity of human annotation from time-lapse images compared to stereomicroscopy.

**D** Visibility of lenses in retinal organoids over time. The dataset was assembled as described in C and annotated by two experts for lens visibility in the respective images. The ground truth was defined by the same two independent researchers and F1-scores were calculated accordingly.  Lenses begin to visibly emerge around 59 h of organoid development.

**E/F** Distribution of RPE (E) and lens (F) classes across experiments. RPE and lens areas were quantified for each RPE and lens positive organoid in the dataset. There was a high variability in class distributions found across experiments, although culture conditions were kept uniform. For the class attributions refer to the Methods and Figure 1C.

### Supplementary Figure S2

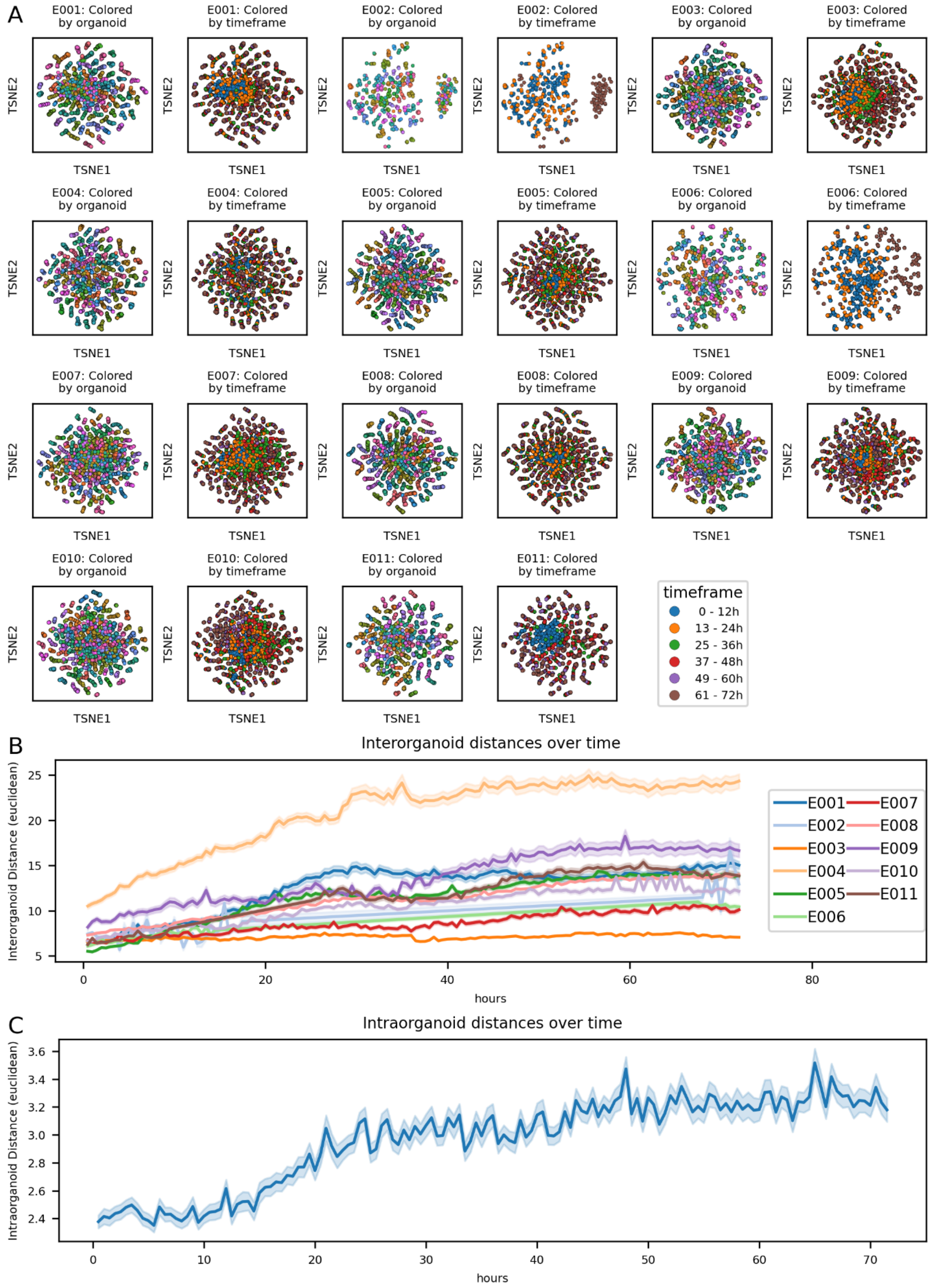

**Supplementary Figure S2: Global morphological heterogeneity of retinal organoids across development between experiments.**

**A** Retinal organoid images were analyzed using the image analysis platform (see Figure 2 and Methods). Data of the indicated experiments (E0XX) were subjected to t-SNE dimensionality reduction and colored by organoid-identity (respective left graph) and the time of acquisition (respective right graph). While the data points of organoid images from earlier time points cluster more closely, there is a substantial divergence of organoids at later time points, suggesting increasing inter-individual differences of the morphologic characteristics over time.

**B** Global morphological inter-organoid heterogeneity across experiments and time. The data correspond to the data shown in Figure 2C. Here, only organoids without Wnt-surrogate treatment were analyzed. Euclidean distances were calculated based on scaled morphometrics data for each organoid and plotted as the mean pairwise distance between all organoids for each time point. Notably, inter-organoid distance increased over time in an experiment-specific manner, indicating an increasing morphological divergence of the individual organoids over time.

**C** Intra-organoid morphological changes over time. The data correspond to the data shown in Figure 2C. Here, only organoids without Wnt-surrogate treatment were analyzed. Euclidean distances were calculated based on the morphometrics data for each organoid between time point *n* and time point *n+1*. The resulting metric reflects the amount of morphological changes during a timespan of 30 min. While the relative changes are comparatively small at the beginning, the increase over time is suggesting more drastic morphological changes at later time points, consistent with the findings described in B and C.

### Supplementary Figure S3

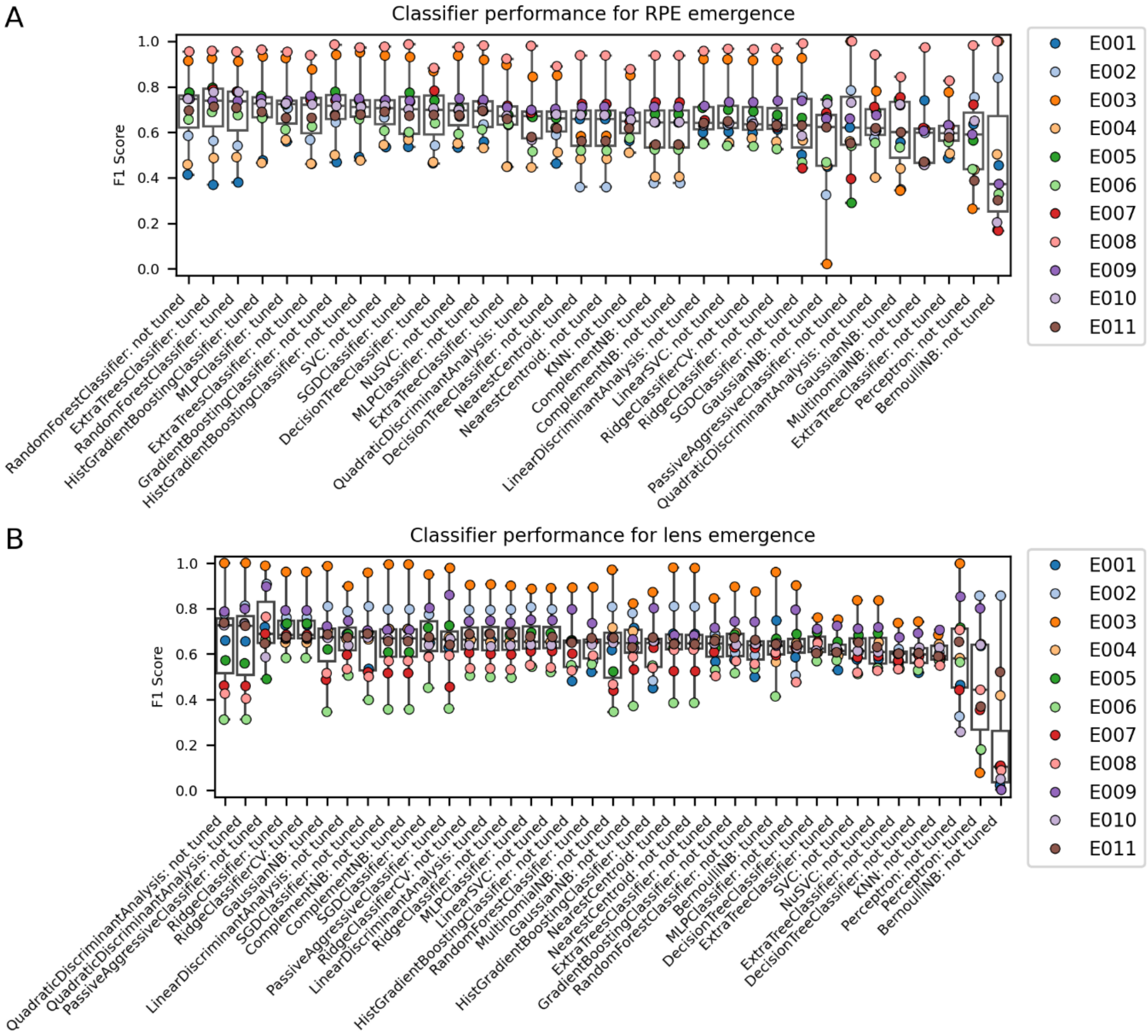


**Supplementary Figure S3: Machine learning classifier benchmark and hyperparameter tuning for tissue emergence predictions.**

**A** The indicated classifiers were trained by cross-validation, using the indicated experiment as a validation set, and scored using the F1 metric (y-axis) for the prediction of the presence and absence of RPE. Selected classifiers were subjected to hyperparameter tuning first (tuned). **B** The indicated classifiers were trained and evaluated as in A, but for the emergence of lenses.

### Supplementary Figure S4

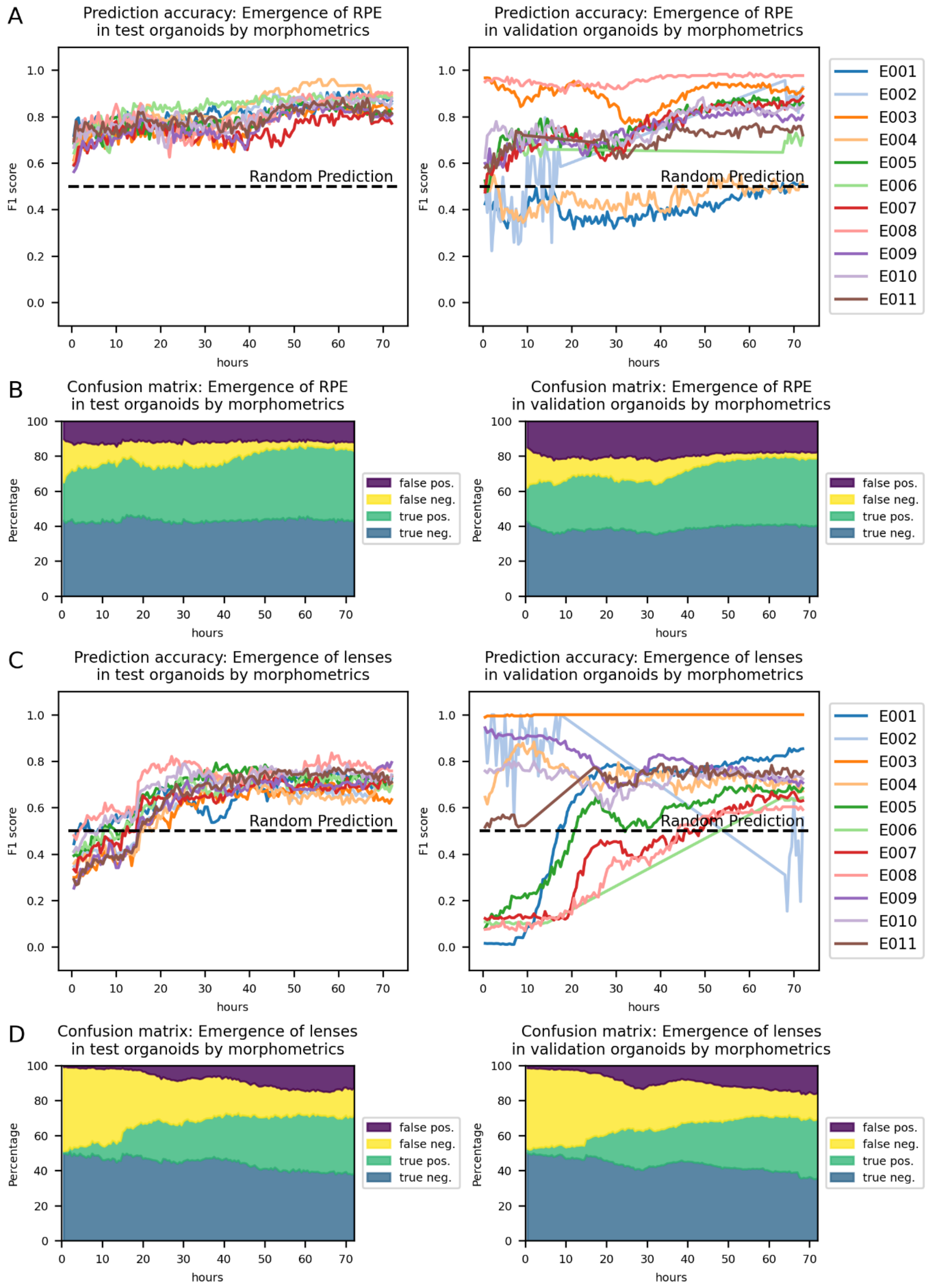

**Supplementary Figure S4: Prediction of tissue emergence by tabular image analysis data.**

Machine learning classifiers were evaluated on the ability to predict RPE emergence (A, B) and lens emergence (C, D) on the test (left graph) and validation (right graph) data sets (for the data partitioning strategy refer to Figure 3A and Methods). **A/C**: The data correspond directly to the data shown in Figure 3B (RPE emergence) and Figure 3C (lens emergence) but are split for the individual experiments. **B/D**: Confusion matrices. The data correspond to A and C, respectively. The x-axis denotes the respective imaging time points while the y-axes show the relative percentage of true-positive, true-negative, false-positive and false-negative predictions as indicated.

### Supplementary Figure S5

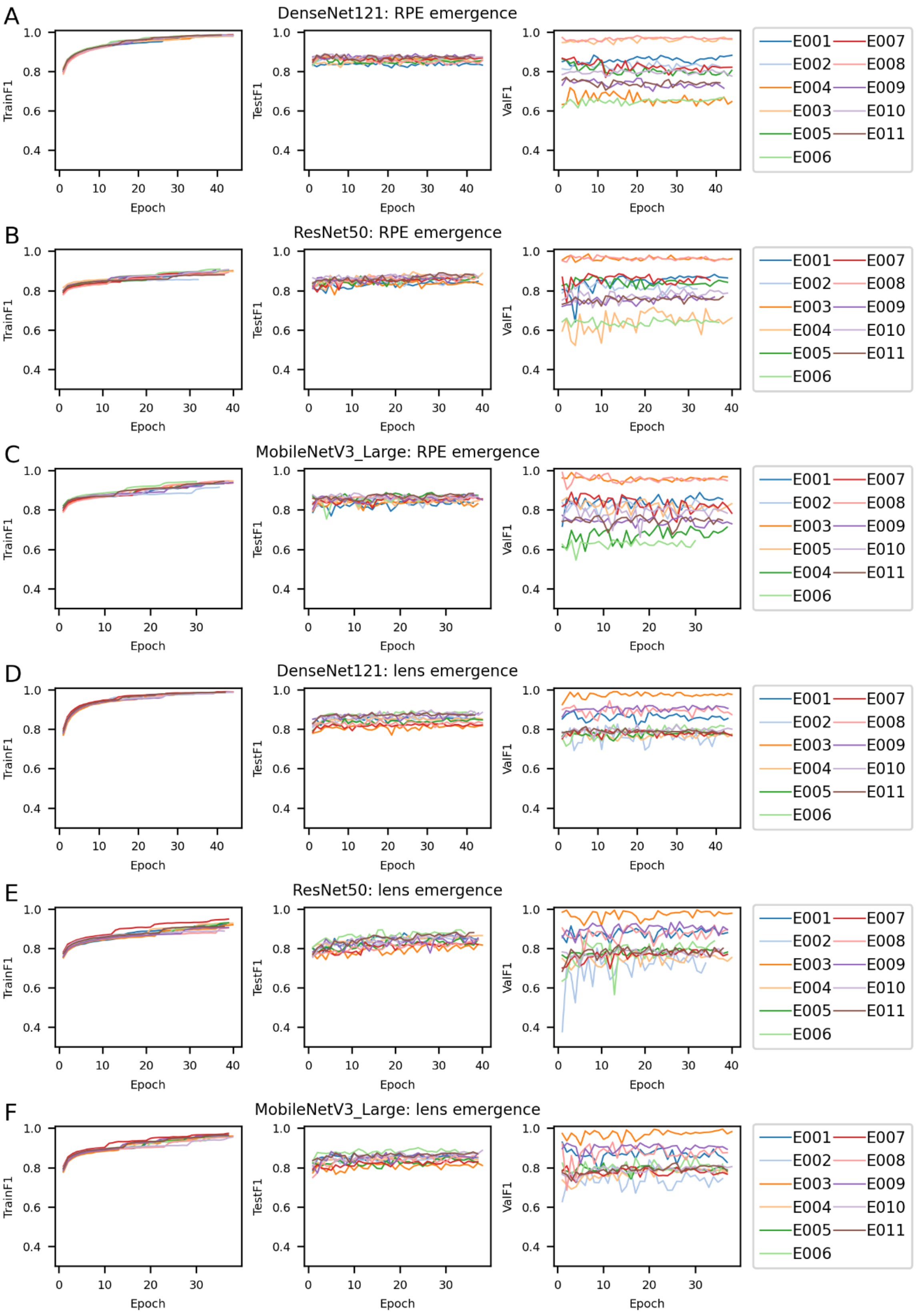

**Supplementary Figure S5: Training and evaluation of CNNs for RPE and lens emergence predictions.**

CNNs were trained to predict the emergence of RPE (**A-C**) and lenses (**D-F**) from time-lapse images for the indicated amount of epochs and scored for the F1-metric in the training set (left graph), the test set (middle graph) and the validation set (right graph). The architecture of the respective CNN is noted within the respective title.

### Supplementary Figure S6

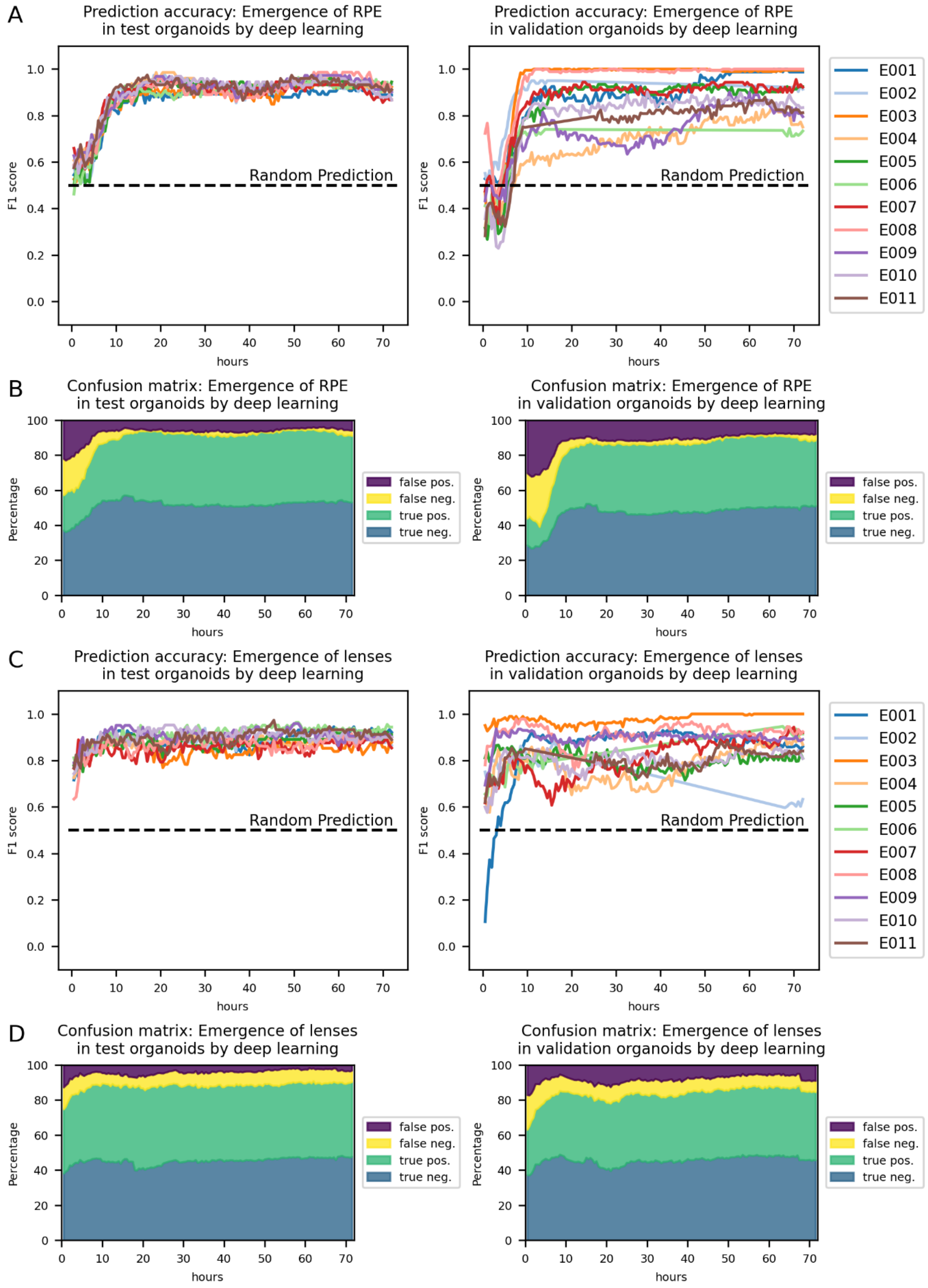

**Supplementary Figure S6: Prediction of tissue emergence by deep learning.**

Deep learning classifiers were evaluated on the ability to predict RPE emergence (A, B) and lens emergence (C, D) on the test (left graph) and validation (right graph) data sets (for the data partitioning strategy refer to Figure 3A and Methods). **A/C**: The data correspond directly to the data shown in Figure 3B (RPE emergence) and Figure 3C (lens emergence) but are split for the individual experiments. **B/D**: Confusion matrices. The data correspond to A and C, respectively. The x-axis denotes the respective imaging time points while the y-axes show the relative percentage of true-positive, true-negative, false-positive and false-negative predictions as indicated.

### Supplementary Figure S7

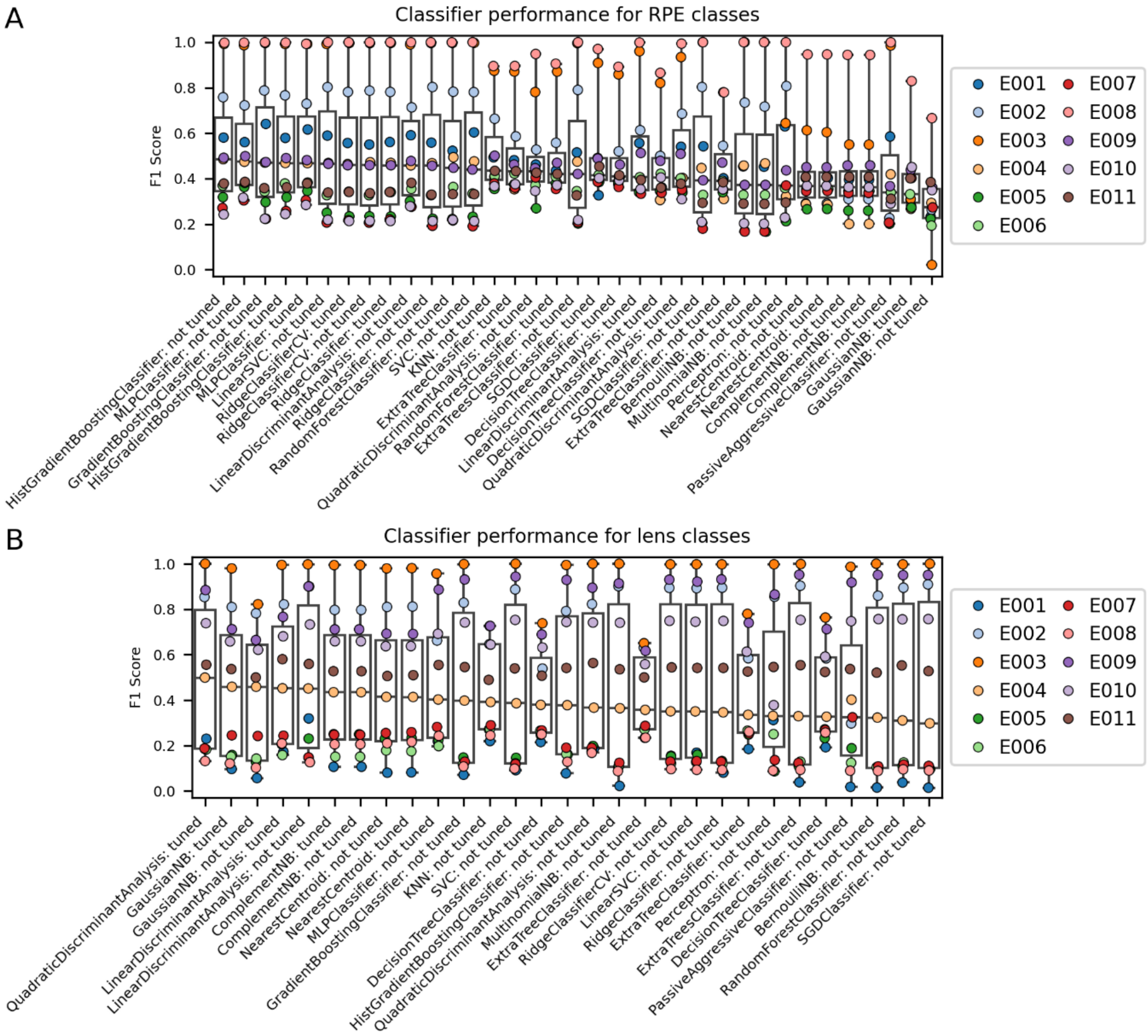


**Supplementary Figure S7: Machine learning classifier benchmark and hyperparameter tuning for tissue size predictions.**

**A** The indicated classifiers were trained by cross-validation, using the indicated experiment as a validation set, and scored using the F1 metric (y-axis) for the prediction of RPE area class. Selected classifiers were subjected to hyperparameter tuning first (tuned).
**B** The indicated classifiers were trained and evaluated as in A, but for the class of lens area.

### Supplementary Figure S8

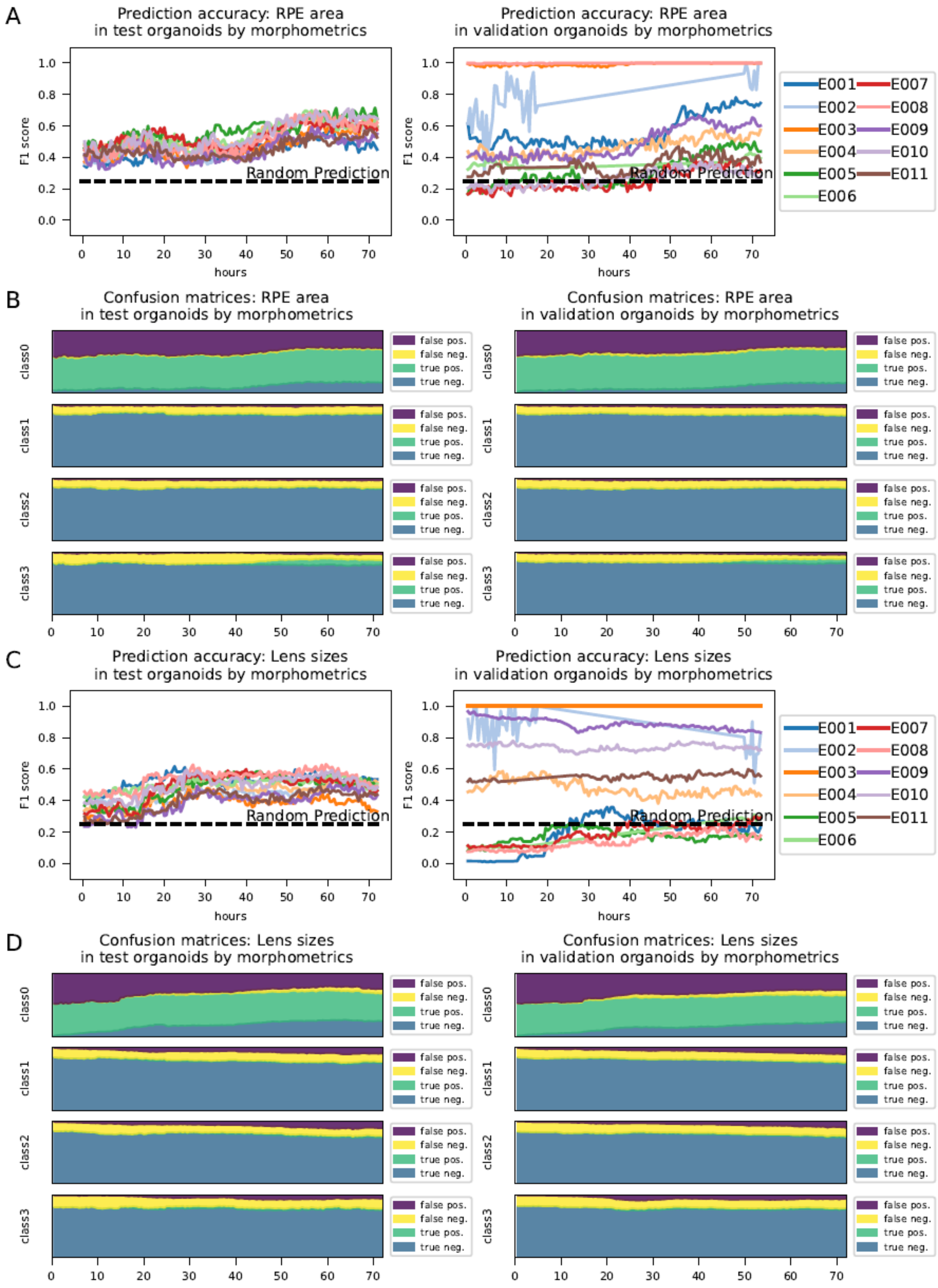

**Supplementary Figure S8: Prediction of tissue sizes by tabular image analysis data.**

Machine learning classifiers were evaluated on the ability to predict RPE areas (A, B) and lens areas (C, D) on the test (left graph) and validation (right graph) data sets (for the data partitioning strategy refer to Figure 3A and Methods). **A/C**: The data correspond directly to the data shown in Figure 4A (RPE emergence) and Figure 4B (lens emergence) but are split for the individual experiments. **B/D**: Confusion matrices. The data correspond to A and C, respectively. The x-axis denotes the respective imaging time points while the y-axes show the relative percentage of true-positive, true-negative, false-positive and false-negative predictions as indicated.

### Supplementary Figure S9

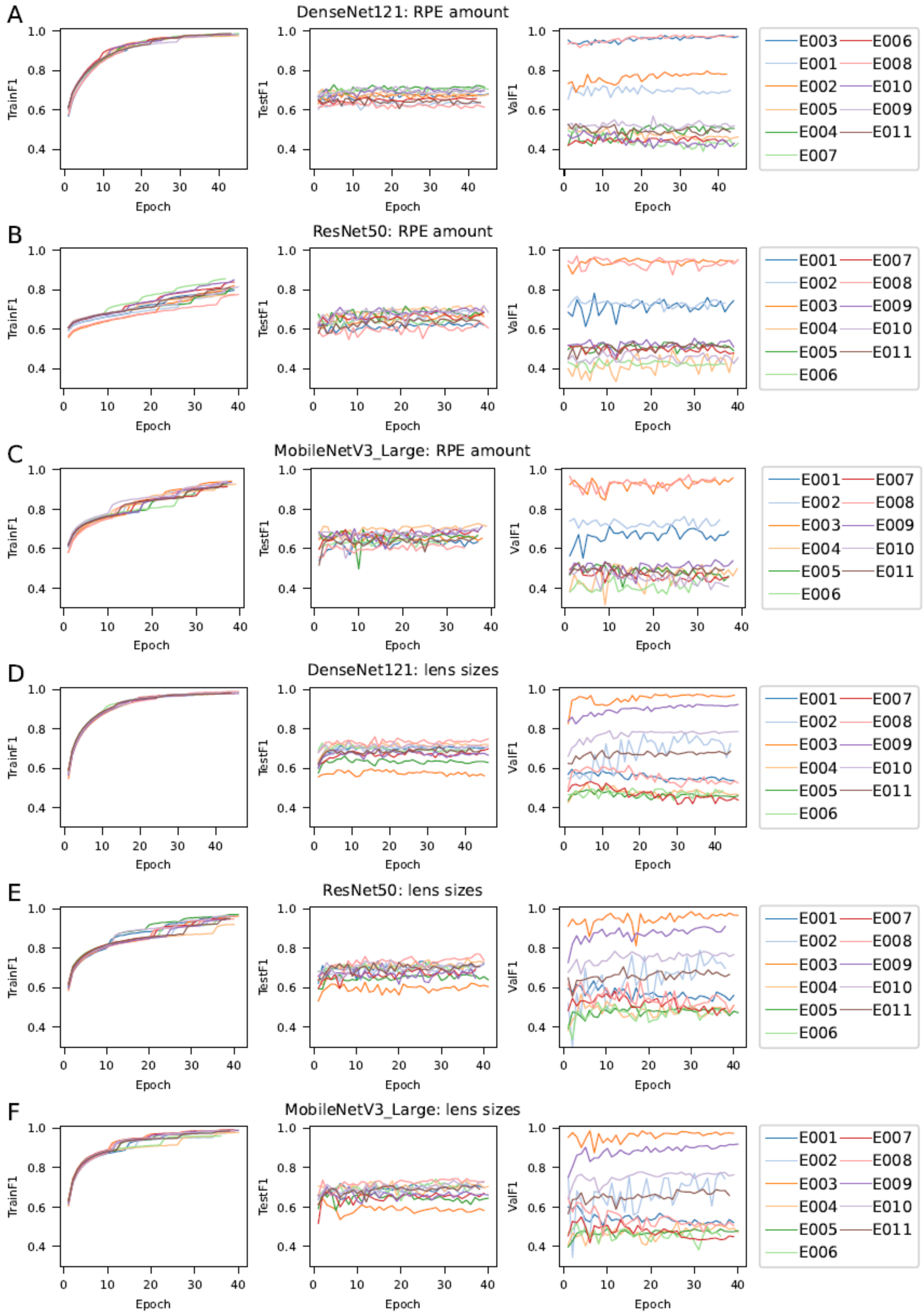

**Supplementary Figure S9: Training and evaluation of CNNs for RPE and lens area predictions.**

CNNs were trained to predict the areas of RPE (**A-C**) and lenses (**D-F**) from time-lapse images for the indicated amount of epochs and scored for the F1 metric in the training set (left graph), the test set (middle graph) and the validation set (right graph). The architecture of the respective CNN is noted within the title.

### Supplementary Figure S10

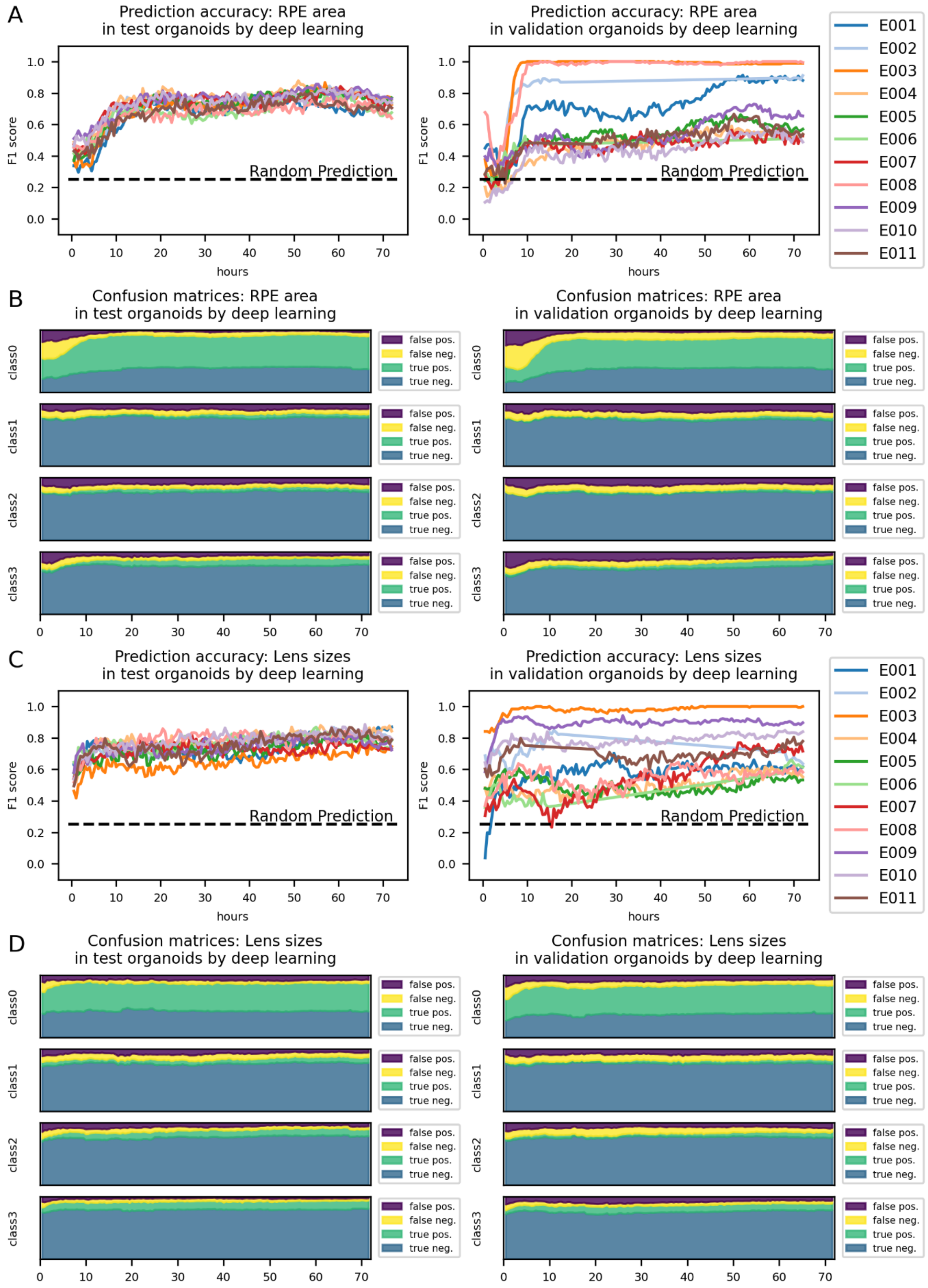

**Supplementary Figure S10: Prediction of tissue size by deep learning.**

Deep learning classifiers were evaluated on the ability to predict RPE areas (A, B) and lens areas (C, D) on the test (left graph) and validation (right graph) data sets (for the data partitioning strategy refer to Figure 3A and Methods). **A/C**: The data correspond directly to the data shown in Figure 4A (RPE emergence) and Figure 4B (lens emergence) but are split for the individual experiments. **B/D**: Confusion matrices. The data correspond to A and C, respectively. The x-axis denotes the respective imaging time points while the y-axes show the relative percentage of true-positive, true-negative, false-positive and false-negative predictions as indicated.

### Supplementary Figure S11

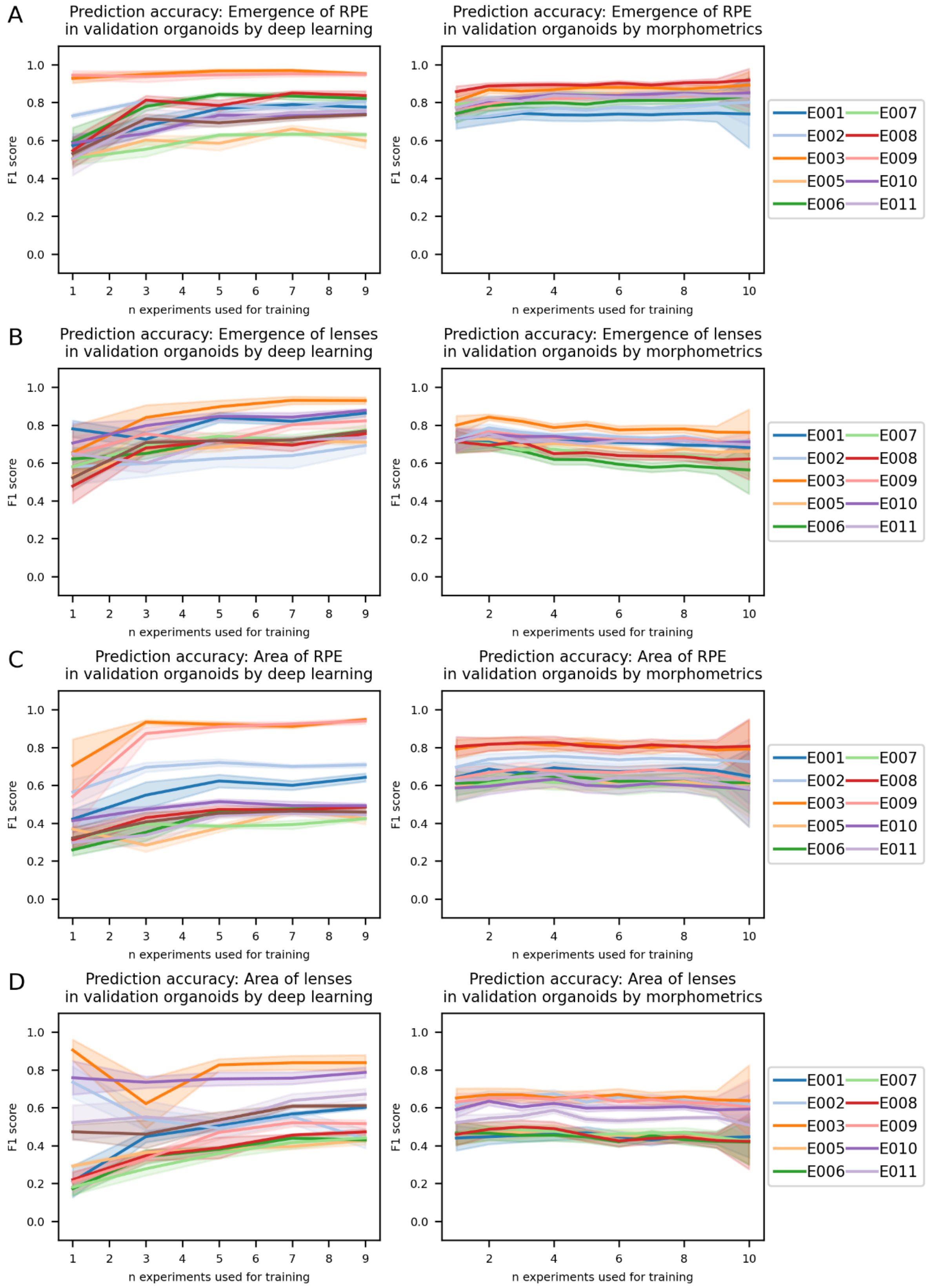

**Supplementary Figure S11: Number of experiments needed for an accurate prediction of TOI.**

Number of experiments needed for an accurate prediction of RPE (**A**) and lens (**B**) emergence, and RPE (**C**) and lens (**D**) area. The experiments corresponding to Figure 3 and Figure 4 were repeated with the indicated number of total experiments (x-axis) used for training. We chose MobileNetV3_Large as the CNN due to computational resources, which was trained for exactly one epoch (left graph), as we did not observe a striking increase in validation accuracy after a higher number of epochs (compare Supplementary Figures 5 and 9). The machine learning classifiers (right graph) were used similarly to Figure 3 and Figure 4, dependent on the classification task. While the curves plateaued at approximately 6 experiments for the classification by neural networks, we could not observe a similar trend for the machine learning classifiers, indicating that tabular data guided classification needs fewer training experiments for a comparable accuracy on the validation set.
